# Sp100-HMG drives ‘inside-out’ PML-NB assembly to modulate transcription and cell-cycle dynamics

**DOI:** 10.1101/2025.07.28.667104

**Authors:** Hongchang Dong, Xiuyan Guo, Yilei Ma, Chaoyi Chen, Jialing Li, Xiao Zhang, Weidong Li, Xiaomei Deng, Honghui Lai, Yaotong Liao, Lin Ye, Pei Xu

## Abstract

Promyelocytic leukemia nuclear bodies (PML NBs) are membraneless organelles (0.1-1 μ m) integral to fundamental cellular processes, yet the molecular logic governing their hierarchical organization is unclear. Contrary to the classical "PML-sole-scaffold" model, we identify Sp100-HMG as the unique isoform that autonomously nucleates into liquid-like condensates. Our findings establish an "inside-out" assembly paradigm, where Sp100-HMG initiates a liquid core that recruits PML and other client proteins (DAXX, ATRX) through three distinct cooperative processes: I. Multimerization domain- and intrinsically disordered region (IDR)-mediated LLPS of Sp100-HMG, nucleating the core; II. C-terminal-dependent protein-protein interactions enriching client components; and III. SUMOylation-directed PML recruitment, facilitating the formation of a stabilizing peripheral shell. This assembly paradigm extends beyond Sp100-HMG, as evidenced by ZBTB16—a PML NB-associated oncoprotein implicated in acute promyelocytic leukemia — adopting an analogous mechanism to organize PML-positive condensates. Using HEp-2 cells as a main model, we show this hierarchical assembly is critical for orchestrating transcriptional programs and cell-cycle dynamics. Together, our study defines a new biogenesis mode for PML-NBs, where a master liquid nucleator coordinates shell integrity and plasticity for nuclear homeostasis.

**Significance Statement:** PML nuclear bodies (PML NBs) serve as essential organizers of the cell nucleus, yet the assembly principles underlying their biogenesis and linking it to functional heterogeneity, spatial positioning, and compositional diversity remain elusive. This study overturns the long-standing "PML-centric" view by establishing an LLPS-driven, hierarchical "inside-out" assembly paradigm. We identify Sp100-HMG as a master liquid nucleator that initiates a core to organize the outer PML shell. This inner-core LLPS-driven assembly logic is shared by other PML NB-associated factors, such as the oncoprotein ZBTB16, providing a unified biophysical framework to explain the structural integrity and plasticity of nuclear condensates. Functionally, this organizational process is critical for shaping the compositional diversity and spatial specificity of PML bodies, thereby modulating transcriptional programs and cell-cycle dynamics in a context-dependent manner. These findings offer broad insights into how organized protein condensation sustains nuclear homeostasis and its dysfunction in diseases such as leukemia.

## Introduction

The eukaryotic nucleus is spatially organized into distinct sub-nuclear compartments known as membrane-less organelles (MLOs), which facilitate the concentration of specific proteins and/or nucleic acids to regulate diverse cellular processes (1). Among these, Promyelocytic Leukemia Nuclear Bodies (PML-NBs) are prominent structures implicated in transcription, apoptosis, DNA damage response, oxidative stress, tumor suppression, chromatin regulation, senescence, and the interferon (IFN) mediated antiviral response (Fan et al., 2023) (2–4). These nuclear condensates dynamically recruit an expanding repertoire of client proteins (5–10). Their functional versatility arises primarily from their ability to function as centers for protein sequestration, modification, degradation, and as platforms for signaling integration and epigenetic control, underpinned by a highly organized and layered structure. Historically, the PML protein (TRIM19) has been regarded as the essential "scaffold" of these bodies. Early genetic studies demonstrated that the loss of PML leads to the dispersion of other nuclear body components, leading to the prevailing model that PML oligomerization—driven by its RBCC (RING finger, B-box1, B-box2, and Coiled-Coil domains) domain —is the primary event in NB biogenesis (11–18). The coiled-coil region of RBCC is rich in intrinsically disordered regions (IDRs) and low-complexity regions (LCRs), which may facilitate multivalent weak interactions (e.g., hydrophobic, electrostatic, π-π stacking) that drive the polymerization of PML protein, and thereby the NBs(19–27). In this classical view, the PML protein serves as the primary scaffold that constitutes the MLO’s outer spherical shell (28). Within this pre-formed or developing PML shell, resident proteins (such as SP100, DAXX, and ATRX) or conditionally associated proteins (such as HIRA, UBC9, RNF4, TDP-43 and p62) are recruited as ’clients’ to populate the inner core (16, 28–37). Both shell formation and client recruitments are reported to be strictly dependent on SUMO-SUMO-interacting motifs (SIMs) interactions (14, 38–41). Manipulation of SUMO1/2/3, SUMO ligases UBC9, SUMO-specific proteases, and within PML or client proteins are reported to strongly disturb the formation of these nuclear bodies (42–47). Notably, PML itself possesses intrinsic SUMO E3 ligase activity, particularly through its B-box1 domain, which facilitates its auto-SUMOylation, promotes LLPS and is believed to initiate nuclear body formation (2, 15, 27, 44, 45, 48–50).

Recent advances have significantly deepened our understanding of PML-NBs. Specifically, the cryo-EM structure of the PML RBCC dimer has illuminated the structural mechanisms underlying PML-NBs assembly, revealing a unique fold and an "octopus-like" oligomerization model.(13, 16, 51, 52). In parallel, concurrent insights into the LLPS characteristics of these MLOs have shed light on their basic principles of spatiotemporal organization (1). Yet, despite these crucial structural and biophysical revelations, a fundamental paradox persists: while pathogenic fusion proteins like PML/RARα may drive condensation, direct evidence for LLPS by PML itself—alone or SUMOylated—remains elusive (53). This complexity is further compounded by the unique architecture of PML-NBs. Characterized by a multilayered organization—specifically, a PML shell surrounding a distinct core—PML-NBs defy categorization as simple liquid condensates, thereby obscuring the underlying biophysical principles that govern their precise assembly and functional regulation (53–60). Moreover, emerging high-resolution analyses further challenge this classical view, revealing that PML can form a shell around pre-existing condensates (e.g., p62) (61, 62). Notably, internal components such as SP100 and SUMO 1/2/3 remain strictly immiscible with the p62 core (63). Collectively, these findings tip the balance of the PML body assembly mechanism toward an "inside-out" assembly mode, where PML proteins/shell is recruited rather than being the primary nucleator. Beyond these structural complexities, a fundamental biophysical challenge persists: the sequence heterogeneity among the seven PML isoforms (PML I-VII) contradicts the remarkable architectural consistency of NBs across cellular contexts. This suggests that the cell utilizes a more evolutionarily conserved molecular switch to impose spatial order onto the variable PML landscape (1, 3, 64). Therefore, deciphering the assembly mechanism of these MLOs, particularly the hierarchical interplay among their dynamic formation/dissolution, SUMO-SIM interaction and LLPS, represents a key step in understanding PML body function and regulation.

Given this requirement for a standardized nucleator, we turned our attention to the Sp100 family. Identified chronologically alongside PML via the analysis of autoantibodies in patients with primary biliary cholangitis (PBC) (65). SP100 represents the first characterized ’client’ of PML nuclear bodies (previously designated ND10 or PODs) (66, 67). It functions as a constitutive resident, localizing to the sub-shell region immediately adjacent to the PML scaffold. Implicated in both intrinsic and innate immunity, SP100 is an Interferon-Stimulated Gene (ISG) that has been reported to support intrinsic defense against HSV-1 infection (68–70). Its functions include modulating ISG expression—potentially via nucleocytoplasmic shuttling—and driving the recruitment of the H3.3 deposition chaperone HIRA to PML-NB (34, 35, 71). Through alternative splicing, the Sp100 gene gives rise to four principal isoforms—Sp100A, Sp100B, Sp100C, and Sp100-HMG—which share a common N-terminal region comprising the first 447 amino acids (70). Despite this shared domain, differences in their C-terminal sequences confer distinct subcellular localizations and biological activities (72, 73). The N-terminal region contains motifs essential for Sp100 dimerization and PML NB targeting (74, 75), including a destruction box (D-box) required for proteasomal degradation, a SUMO consensus motif, a SUMO-interacting motif (SIM) within a histone protein 1 (HP1) binding site, and a transactivation region (74, 76–78). Owing to its multifaceted functionality, dysregulation of Sp100 has been implicated in diverse pathological contexts, including oncogenesis and viral pathogenesis (69, 71). As a prominent PML NB-associated factor, Sp100 is widely regarded as a core client protein contributing to the assembly, signaling, and maintenance of these nuclear compartments. Nevertheless, the precise molecular mechanisms by which Sp100 participates in PML NB biogenesis and exerts its functional effects through these structures remain poorly understood.

An initial observation that Sp100-HMG undergoes LLPS by itself prompted a systematic investigation into its role in PML-NB biogenesis in this report. We delineate a stepwise assembly model governed by phase separation: (I) Condensate core formation driven by Sp100-HMG’s LLPS mediated by its multimerization domain and intrinsically disordered region (IDR); (II) Recruitment of client proteins (e.g., DAXX and ATRX) via C-terminal-dependent interactions; and (III) Subsequent formation of the peripheral PML shell through SUMOylation-dependent recruitment. A comparable mechanism was observed for the APL-linked protein ZBTB16 (79–81). Collectively, these findings suggest a distinct Sp100-HMG-driven assembly pathway that diverges from the canonical PML-centric model, redefining the role of PML polymerization as a secondary encapsulation step rather than the primary organizing event under these experimental settings. Furthermore, by elucidating the mechanisms of transcriptional and cell cycle regulation by an artificial-core organized PML positive condensates and Sp100-HMG positive PML NBs, this study reveals compelling paradigms of PML NB functional regulation that are inherently tied to their unique internal physical architecture.

## Results

### LLPS of Sp100-HMG leads to de novo formation of PML NBs

While PML is the hallmark component of NBs, the lack of evidence for its spontaneous LLPS suggests the existence of other organizing nucleators (33, 37, 82, 83). Given the shell-core architecture of PML-NBs, we hypothesized that internal core components drive assembly. By systematically screening the four major Sp100 isoforms, a constitutive resident protein family of the NB core, in Sp100-/- and PML/Sp100-/- (double knock out, DKO) HEp-2 cell lines (68, 83), we identified Sp100-HMG as the unique isoform capable of autonomous nucleation into spherical, liquid-like nuclear condensates that exhibited dynamic fusion and fission in the absence of other NB scaffolds (Fig. 1 A, B, Fig. S 1 A).

**Fig. 1.**
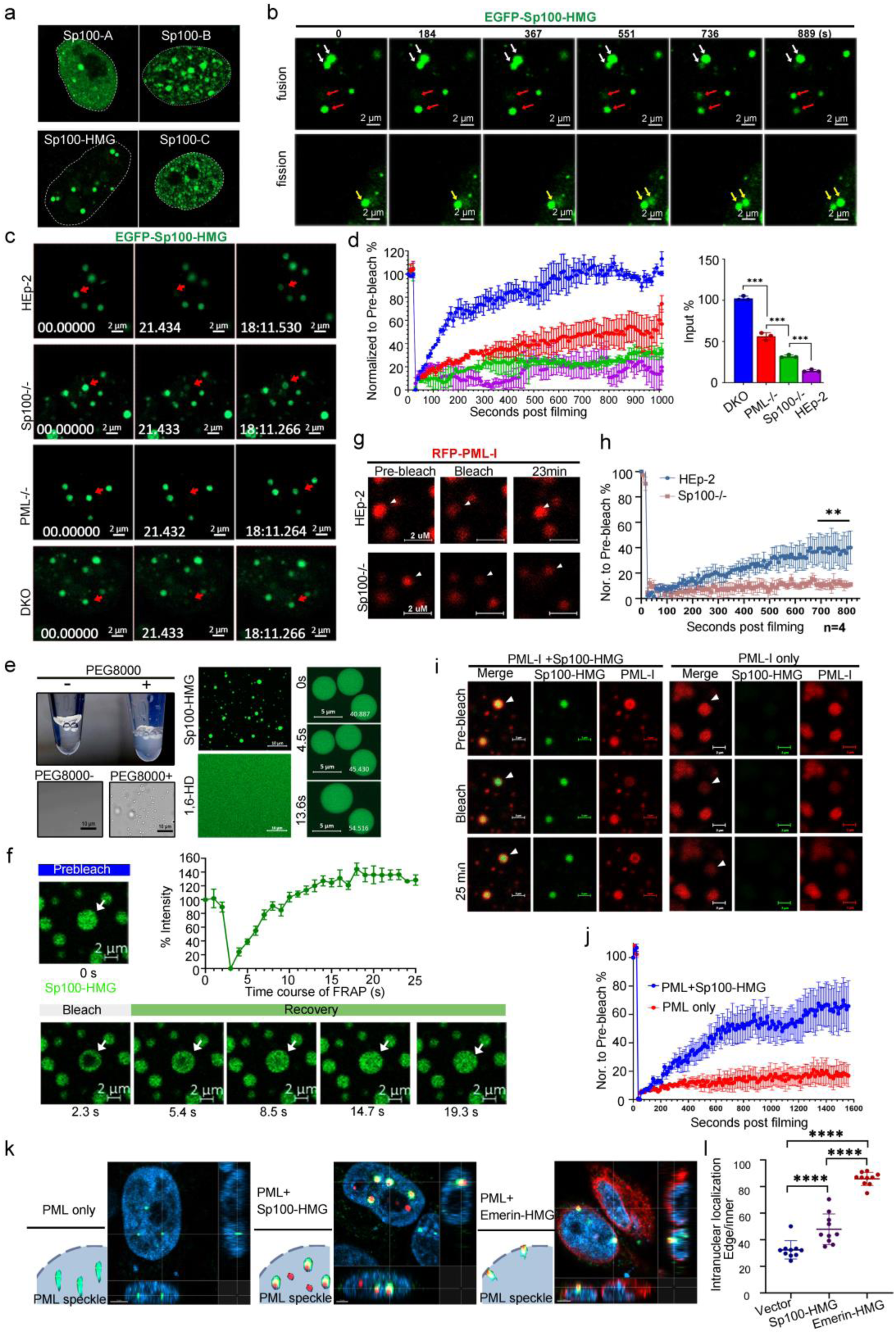
Sp100-HMG forms LLPS in the cell nucleus that recruits PML protein. A. Flag-tagged Sp100-A, -B, -C, and -HMG were transfected into DKO cells for 24 hr and stained with anti-Flag antibody. B GFP-Sp100-HMG was transfected into DKO cells for 12 hr and time-lapse images were taken at the indicated time, revealing fusion and fission of GFP-Sp100-HMG condensates as indicated by arrows. C-D. GFP-Sp100-HMG was transfected into the indicated cells. At 24 hr post-transfection, a defined region (red arrows) within a GFP-Sp100-HMG condensate was subjected to FRAP. Representative time-lapse images for each cell line are shown in C and corresponding fluorescence recovery kinetics are quantified in D. E. Bacterially purified full-length human Sp100-HMG was labelled with FITC. Turbidity (up) and bright filed images (down) of Sp100-HMG solution with or without PEG8000 (LLPS conditions: Sp100-HMG 12.5μM, Nacl 100μM, 15% m/v% PEG8000) (Left panel). Fluorescence images of Sp100-HMG condensates with or without 10% 1,6 HD treatment (LLPS conditions: 15% PEG8000, Sp100-HMG 50 μM, Nacl 100μM) (Middle panel). Time-lapse images of the merging Sp100-HMG droplets (Right panel). F. FRAP of in vitro Sp100-HMG condensates. Left: Representative time-lapse images before and after photobleaching. Right: Fluorescence recovery curves of Sp100-HMG within the droplets. G-H. FRAP of transfected mCherry-PML in HEp-2 vs. Sp100-/- cells. G. Representative time-lapse images; arrows indicate bleached spots. H. PML recovery kinetics. I-J. FRAP of mCherry-PML in DKO cells w/wo GFP-Sp100-HMG. I. Representative images showing mCherry-PML and GFP-Sp100-HMG recovery (arrows: bleached spots). J. PML recovery kinetics. K. Representative images and illustrations of endogenous PML in Sp100-/- cells transfected with Sp100-HMG or Sp100-HMG-Emerin. Peripheral localization is defined as adjacency to the nuclear envelope. L. Quantification of peripheral PML speckles in K per group.

Fluorescence recovery after photobleaching (FRAP) analysis demonstrated that Sp100-HMG dynamics are physically constrained by the NB architecture: its recovery rate was minimal in wild type HEp-2 cells but increased progressively upon the loss of endogenous Sp100 and PML, peaking in DKO cells (Fig. 1 C, D). This step-wise increase suggests that the PML-NB scaffold imposes a hierarchical restriction on the mobility of the internal Sp100 core. To validate these cellular findings, we reconstituted the system in vitro. Bacterially purified Sp100-HMG underwent LLPS at a physiologically relevant concentration of 12.5 μM upon molecular crowding, forming highly fluid, 1,6-HD-sensitive droplets (Fig. 1 E, F) (84–86). Detailed phase-mapping showed that Sp100-HMG nucleation is driven by electrostatic interactions, as propensity increased with decreased ionic strength and neutral pH (Fig. S 1 B-F).

We next challenged the paradigm of PML as the sole essential scaffold by introducing Sp100-HMG into NT2 cells (11). Ectopic expression of Sp100-HMG in NT2 cells significantly augmented the volume of existing NBs, forming a core that efficiently recruited classical clients like DAXX and ATRX (Fig. S 1 G-I). Notably, this Sp100-HMG/PML interplays in NT2 cells recapitulated the reported p62-PML recruitment patterns (Fig. S 1 G, H), suggesting a conserved ’inside-out’ assembly principle where an immiscible liquid core organizes the outer PML shell (63).

Furthermore, while PML NBs can persist in the absence of Sp100, their marked reduction in size and abundance points to a structural collapse or instability of the PML shell when deprived of its internal liquid scaffold (Fig. S1 G, J-L) (72). Our biophysical analysis further revealed a striking interplay between these layers: while the PML-I shell is relatively rigid, the presence of the dynamic Sp100-HMG core actively fluidized the PML-I shell, increasing its exchange rate in both HEp-2 and NT2 cells (Fig. 1 G-J, Fig. S 1 M, N).

Finally, using a nuclear membrane-tethering assay, we established the hierarchical directionality of the assembly of PML NBs. Emerin-tagged Sp100-HMG successfully nucleated *de novo* PML-positive puncta at the nuclear periphery in both HEp-2 and non-cancerous HaCaT cells, whereas Emerin-tagged PML-I failed to recruit Sp100-HMG (Fig. 1 K, L, Fig. S 1 O, P). Notably, Sp100-HMG overexpression shifted PML NBs toward the nuclear periphery (Fig. 1 I, Fig. S 1 O), likely driven by a synergy between HMG-heterochromatin targeting and the biophysical exclusion of large condensates. Collectively, these results demonstrate that Sp100-HMG serves as a dynamic internal scaffold that not only directs PML assembly but also fluidizes the outer PML shell that initiates PML NBs formation and imposes liquid-like properties onto the outer PML shell via an "inside-out" assembly mode.

### Sp100-HMG organizes PML NBs via domain-specific phase separation and SUMO-dependent recruitment of PML

To delineate the molecular domains underpinning Sp100-HMG LLPS, we utilized the IUPred2A algorithm to identify 10 putative disordered segments and 5 are located at the unique C-terminal of Sp100-HMG (Fig. S 2 A) (87). Systemic mutagenesis revealed that segments Dis8, Dis9, and Dis10 are indispensable for condensation; their replacement with flexible linker-GGSGGS (ΔDis8, ΔDis9, ΔDis10) completely abrogated condensate formation and the subsequent recruitment of DAXX and ATRX (Fig 2 A, Fig. S 3 A). Notably, single-residue screening identified I645 within Dis8 as a critical node for phase separation (Fig. S 3 B). Beyond the C-terminal IDRs, N-terminal multimerization proved equally vital, as disruption of the polymerization domain (residues 3–152) (Mut 152) or the HP1-binding domain (Mut HP1) severely compromised LLPS (Fig. S 2 B, Fig. S 3 C-E). However, a minimal construct comprising the N-terminal polymerization domain fused to the C-terminal tail (152-cT) was sufficient to form spherical, 1,6-HD-sensitive liquid condensates in DKO cells (Fig. 2 B, C).

**Fig. 2.**
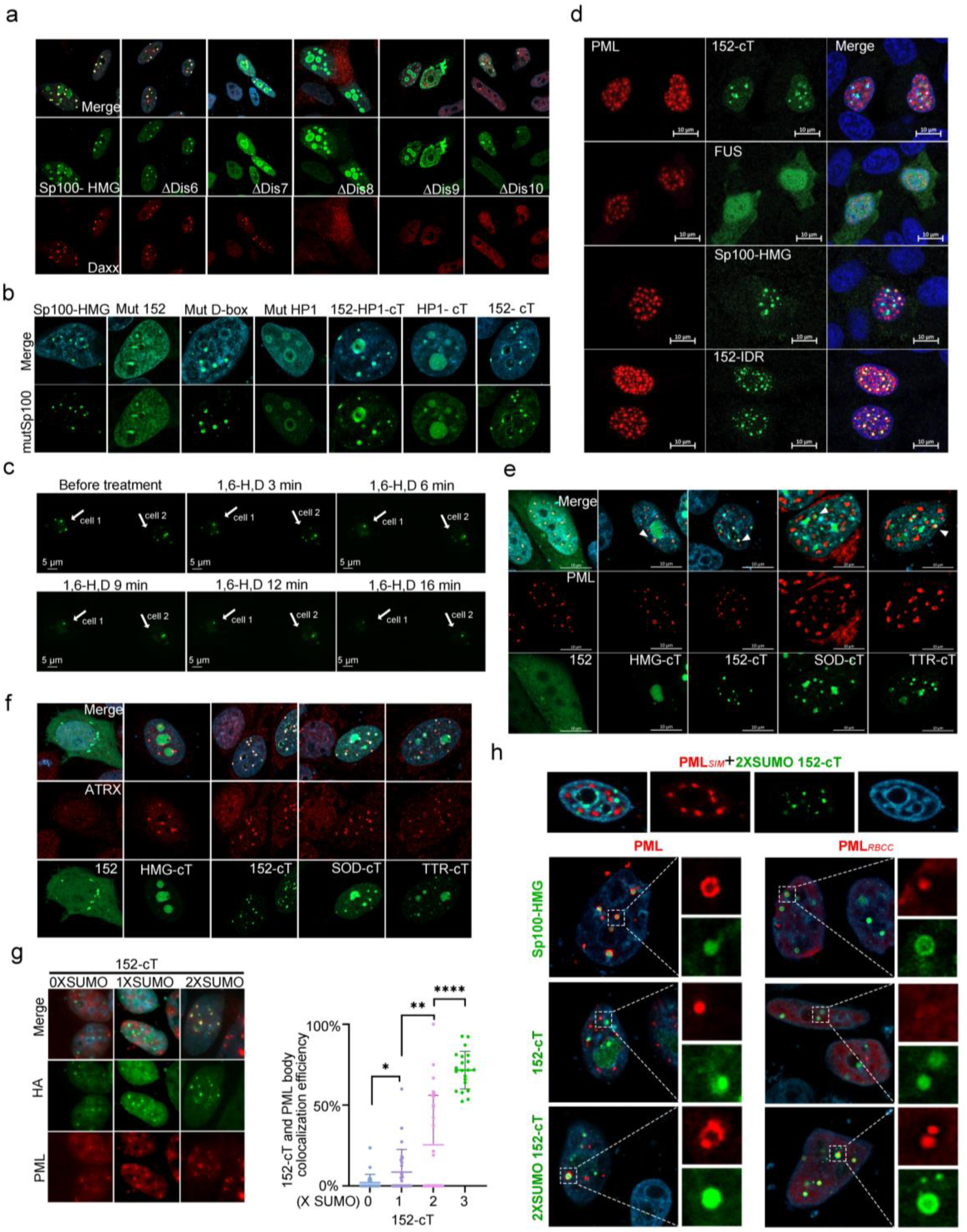
Domain mapping and molecular mechanisms underlying Sp100-HMG–mediated LLPS and recruitment of PML or client protein ATRX. A-B. Screening of C-terminal IDRs (A) (Green: Flag-Sp100-HMG and its mutants and Red: endogenous DAXX) and N-terminal domains (B) mediating LLPS in DKO cells via immunofluorescence staining (Green: Flag-Sp100-HMG and its mutants). C. Representative images of GFP-152-cT condensates in DKO cells treated with 10% 1,6-HD. D. Expression patterns of GFP tagged 152-cT, FUS, Sp100-HMG or 152-IDR fusions co-transfected with PML-I in DKO cells. E. Evaluation of the polymerization function of 152 domain by replacement with tetramerization (TTR) or dimerization (SOD1) domains, cells were co-stained with anti-PML antibody. F. ATRX recruitment by 152-cT and TTR/SOD1-cT variants in DKO cells. G. Right panel, PML recruitment by 152-cT fused with indicated SUMO repeats. Left panel, Statistical quantification of colocalization efficiency. H. Recruitment of PML*_RBCC_* or PML*_SIM_* by HMG, 152-cT, or 2XSUMO-152-cT scaffolds in DKO cells.

To delineate the biophysical contributions of individual domains, we engineered domain-swapped chimeras: I.152-IDR: The C-terminal tail of Sp100-HMG (cT) was replaced with the FUS IDR (152-IDR) (25); II. SOD-cT & TTR-cT: The N-terminal 3-152 sequence was substituted with dimerization (SOD) or tetramerization (TTR) modules (88, 89). While all chimeras formed dynamic nuclear condensates and recruited endogenous PML, their scaffolding capacities were distinct (Fig 2 D, E, Fig S 3 F). Notably, constructs containing the native Sp100-HMG C-terminal tail efficiently recruited ATRX (Fig 2 F). This allowed us to observe functional decoupling between shell organization and client sequestration. The 152-IDR chimera efficiently organized a prominent PML shell but failed to sequester DAXX. In sharp contrast, the 152-cT construct efficiently recruited ATRX despite compromised shell formation (Fig. 2 D-F; Fig. S 3 G). These findings establish the Sp100-HMG core, rather than the PML shell, as the autonomous driver of "inside-out" assembly, decoupling internal protein sequestration from the peripheral architecture. To evaluate scaffolding specificity, we examined recruitment of the stress-responsive STUbL, RNF4 to PML NBs (19, 90). While Sp100 depletion abolished IFN-induced RNF4 recruitment, ectopic Sp100-HMG expression was sufficient for its sequestration even without stimulation (Fig S 3 H-J). Importantly, RNF4 showed efficient recruitment at low levels but remained nucleoplasmic as concentrations rose, indicating saturable, specific partitioning rather than an overexpression artifact (Fig S 3 J). Notably, RNF4 exhibited a higher recruitment threshold than ATRX. While 152-cT recruited ATRX, RNF4 sequestration required either the full-length protein or the SUMO-152-cT variant (Fig S 3 K). This suggests that RNF4 recruitment relies on synergy between the C-terminal scaffold and the high-density multivalent SUMO environment provided by the native N-terminal domain. Critically, we observed a functional decoupling between shell organization and client recruitment.

Furthermore, the 152-cT construct exhibited inefficient PML recruitment compared to 152-IDR, suggesting that the molecular requirements for shell organization are distinct from those governing core formation (Fig 2 D, E). Given that SUMO-SIM interactions are established determinants for NB stoichiometry (91, 92), we investigated whether increasing SUMOylation or numbers of SIM motifs could rescue PML recruitment to the 152-cT core. Quantitative analysis revealed a dose-dependent enhancement of PML colocalization correlating with the number of SUMO tags, whereas increasing the number of SIM motifs on the Sp100 core had no effect (Fig. 2 G, Fig. S 3 I). This recruitment was strictly mediated by the PML SIM domain, as its genetic ablation abolished recruitment by the 2XSUMO-152-cT variant (Fig. 2 H).

Previous studies have established the RBCC domain as essential for de novo PML nuclear body formation, with arsenic trioxide binding to this region inducing a gel-like transition of PML NBs (16, 19, 20, 51). We found that while a PML mutant lacking the RBCC domain (PML*_RBCC_*) could still be recruited by Sp100-HMG and 2XSUMO-152-cT via SUMO-SIM interactions, unlike the recruitment of the PML mutant with lacking the SIM motif (PML*_SIM_*), it failed to organize into a peripheral shell. Instead, PML*_RBCC_* accumulated as static clusters within the internal core of the condensates (Fig. 2 H). FRAP kinetics confirmed that PML*_RBCC_* was highly immobile, with recovery rates comparable to static protein aggregates, whether existing independently or within the Sp100-HMG matrix (Fig. S 3 M). These results indicate that while SUMO-SIM interactions are sufficient for the initial docking of PML to the scaffold, RBCC-dependent multimerization is indispensable for the subsequent fluid-like organization of the peripheral shell (19).

Collectively, these findings establish a hierarchical ’inside-out’ model of PML NB biogenesis, where Sp100-HMG initiates a liquid core driven by N-terminal multimerization and C-terminal IDRs that subsequently recruits PML via SUMO-SIM docking, while the RBCC domain of PML is essential for the spatial realization of the ’inside-out’ assembly model; it provides the structural integrity required for shell maturation and prevents recruited PML from collapsing into the Sp100-HMG core.

### PML bodies induced by LLPS of SP100-HMG strengthens local transcriptional regulation

To validate the generalizability of our proposed model, we performed a targeted evaluation of high-confidence candidates identified from the literature that were potentially capable of nucleating PML-positive condensates. While the following three criteria could theoretically apply to a broad range of proteins, we focused on a representative cohort of five candidates to provide a proof-of-principle for our ’inside-out’ assembly logic: (I) documented association with PML-NBs and the presence of SUMOylation sites; (II) the presence of IDRs; and (III) a structured multimerization domain. We anticipate that other proteins sharing this modular architecture may follow similar nucleation principles. Among the screened candidates, ZBTB16 (also known as PLZF) emerged as a potent nucleator, forming distinct nuclear puncta even in PML-/- cells (Fig. S 4 A, Fig. 3A), thereby supporting the model that specific multivalent scaffolds can autonomously drive the assembly of the PML NB core. Next, structure-guided mutagenesis revealed that both the BTB/POZ multimerization domain and the predicted IDR (aa 126–189) are essential for phase separation. Disruption of these domains (mutants Mut*_BTB_*, Mut*_100-200_*, and Mut*_100-300_*: deletion of both IDRs and the RD2 domain) resulted in a diffuse nuclear distribution, regardless of the presence (HEp-2 cells) or absence (PML-/- cells) of endogenous PML (Fig. 3 B, C). Notably, while a SUMOylation-deficient mutant (Mut*_3KR_*) retained the ability to form condensates, it failed to recruit PML, confirming that PML recruitment is a secondary, SUMO-dependent "docking" step, functionally decoupled from the initial nucleation event (Fig. 3 C). Interestingly, ZBTB16 condensates selectively recruited DAXX but not ATRX more efficiently, suggesting that the specific identity of the liquid nucleator dictates client composition (Fig. S 4 B, C). These results demonstrate that ZBTB16, like Sp100-HMG, functions as an alternative scaffold that initiates PML-NB assembly through an LLPS-driven "inside-out" mechanism, providing a biophysical basis for the compositional heterogeneity observed in endogenous nuclear bodies.

**Fig. 3.**
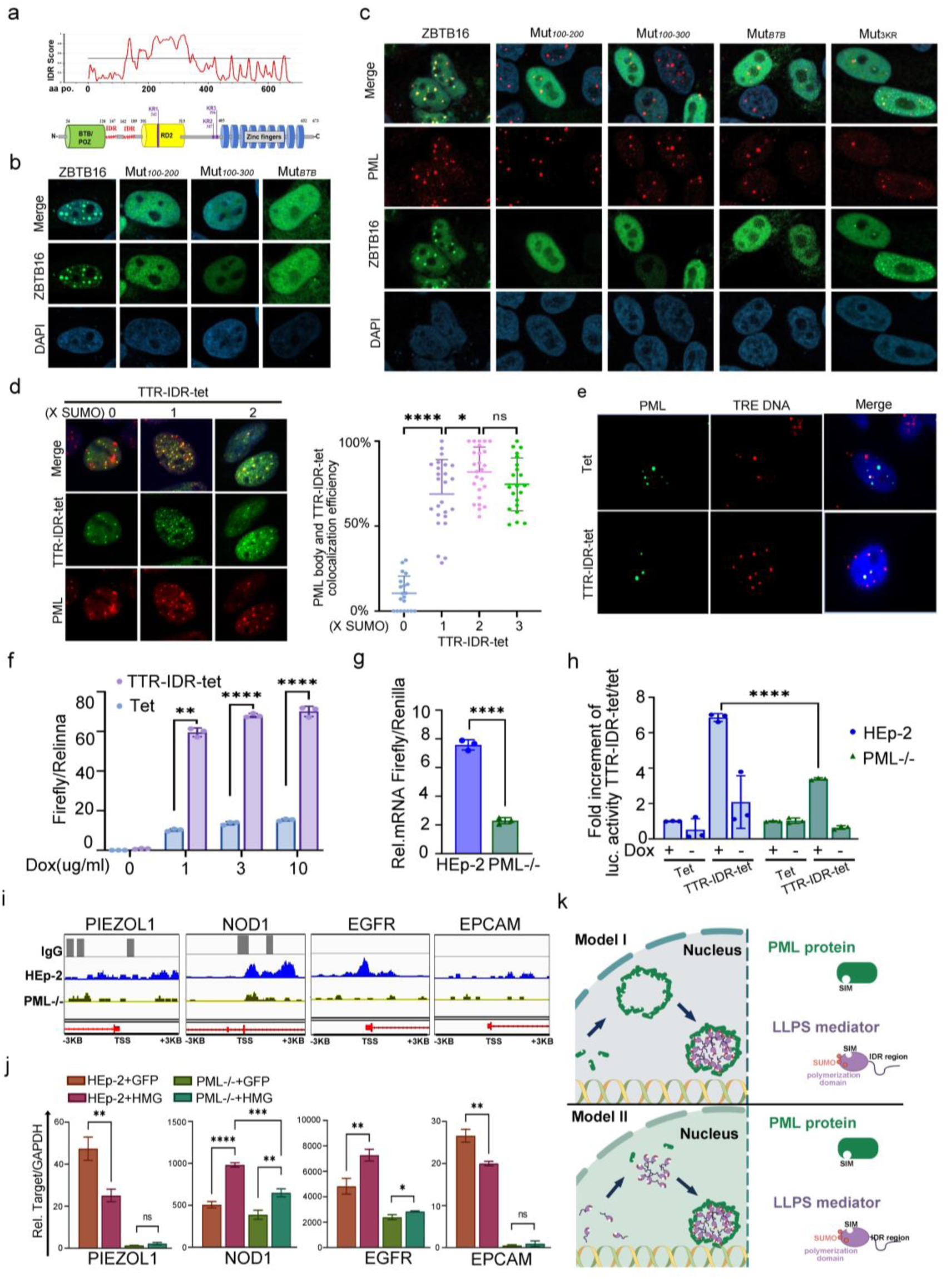
A model for PML NBs *de novo* formation and their role in gene transcription regulation. A. Schematic of ZBTB16 domains and IUPred3.0-predicted IDRs. B. Intranuclear localization of GFP-ZBTB16 and deletion mutants (aa100-200 deletion mutant: Mut*_100-200_*, aa100-300 deletion mutant: Mut*_100-300_*, BTB region deletion mutant: Mut*_BTB_*) in DKO cells. C. Colocalization of GFP-ZBTB16 or Mut*_3KR_* (all 3 lysines indicated in A to arginine) with endogenous PML in DKO cells. D. Artificial PML organizer recruitment. TTR-IDR-tet with indicated number of SUMO repeats in Sp100-/- cells. Left, representative images. Right, colocalization efficiency of TTR-IDR variants with PML (A total 53, 57, 27, 78 cells in each corresponding group were calculated). E. DNA FISH (TRE probe) and PML IF in HEp-2 cells transfected with TRE-plasmids with Tet or TTR-IDR-tet and treated with Dox. F-H. Luciferase activity (F, H) or mRNA levels via qRT-PCR (G) in HEp-2 or PML-/- cells transfected with a Tet-On plasmid expressing TRE-Firefly luciferase and Tet or TTR-IDR-tet treated with indicated Dox concentrations (F) or at 1 ug/ml (G, H) for 48 hr. I. Cut&Tag analysis showing Sp100-HMG enrichment at candidate promoters in HEp-2 and PML-/- cells. J. mRNA levels (qRT-PCR) of the indicated genes in HEp-2 and PML-/- cells in the presence of GFP or Sp100-HMG expression, normalized to GAPDH. K. Proposed assembly models. Schematic illustrating two hypothetical mechanisms for PML NB assembly.

To interrogate the transcriptional regulatory potential of Sp100-HMG-mediated PML nuclear body formation, we engineered a synthetic transcriptional regulator that mimics the biophysical properties of Sp100-HMG. This construct (TTR-IDR-tet) comprises a reverse tetracycline-controlled transactivator (rtTA) fused to an IDR and a tetramerization domain. TTR-IDR-tet formed nuclear condensates that recruited endogenous PML in a SUMO-dependent manner (Fig. 3 D). In HEp-2 cells, TTR-IDR-tet effectively recruited endogenous PML positive puncta formation to tetracycline response element (TRE)-containing loci, significantly enhancing doxycycline-dependent luciferase expression (Fig. 3 E, F). This transcriptional activation was markedly attenuated inPML-/- cells, indicating that the PML is indispensable for the full functional capacity of the condensate (Fig. 3 G, H) The critical dependence on PML was demonstrated by the marked attenuation of this transcriptional enhancement in PML-deficient cells, suggesting that PML shell formation potentiates the transcriptional activation capacity of TTR-IDR-tet condensates (Fig. 3 G-H). These data suggest that the PML shell serves as a functional potentiator for transcriptional regulation.

We further examined the endogenous transcriptional regulatory function of Sp100-HMG using genome-wide mapping via Cut&Tag sequencing (Fig. S 4 D). Sp100-HMG exhibited preferential association with transcriptional start sites (TSS), a pattern that was significantly disrupted in PML-/-, Sp100-/- cells or DKO cells, highlighting a requirement for intact NB architecture (Fig. S 4 E, Fig 3 I). While Sp100-HMG overexpression modulated the expression of target genes (e.g., *NOD1*, *EGFR*, *PIEZOL1*, *EPCAM*) in wild-type cells, these regulatory effects were substantially neutralized in the absence of PML (Fig. 3 I, J). Note that, two of six selected target genes showing PML-dependent Sp100-HMG promoter enrichment failed to exhibit transcriptional responses to Sp100-HMG overexpression, suggesting possible additional layers of regulation (Fig. S 4 F). These findings support a model (Model II) in which Sp100-HMG acts as a master nucleation factor that directs PML-NB assembly to specific genomic loci. In this framework, the hierarchical organization of the NB modulates local transcriptional activity by spatially concentrating or sequestering regulatory factors (Fig. 3 K).

### Sp100-HMG positive PML NBs regulates cell cycle progression by modulating the nucleoplasm availability of DAXX

Given their unusually large dimensions, PML NBs are thought to function as dynamic hubs for the reversible sequestration of proteins and the coordination of post-translational modifications in response to cellular signaling. We first investigated the organizational logic of Sp100-HMG-positive NBs. Our previous observations in this report established that the recruitment of DAXX and ATRX to PML NBs is strictly dependent on Sp100 expression (Fig S 1 I, Fig 2, Fig 4A, Fig S 5 A, B), and that Sp100-HMG is sufficient to restore the punctate organization of these proteins in Sp100-deficient cells (Fig 4 B). To probe the internal dynamics of these structures, we performed FRAP. In PML-/- cells, DAXX exhibited high mobility with nearly complete recovery. However, in HEp-2 cells, DAXX recovery was significantly restricted, plateauing at approximately 60% (Fig 4 C, D). Notably, Sp100-HMG displayed even more constrained dynamics, showing immediate and pronounced immobility compared to DAXX. This gradient of decreasing mobility—from the interior to the periphery—reveals a concentric stratification of the nuclear body architecture, where the outer PML shell imposes physical and molecular constraints on the internal Sp100-DAXX scaffold.

**Fig. 4.**
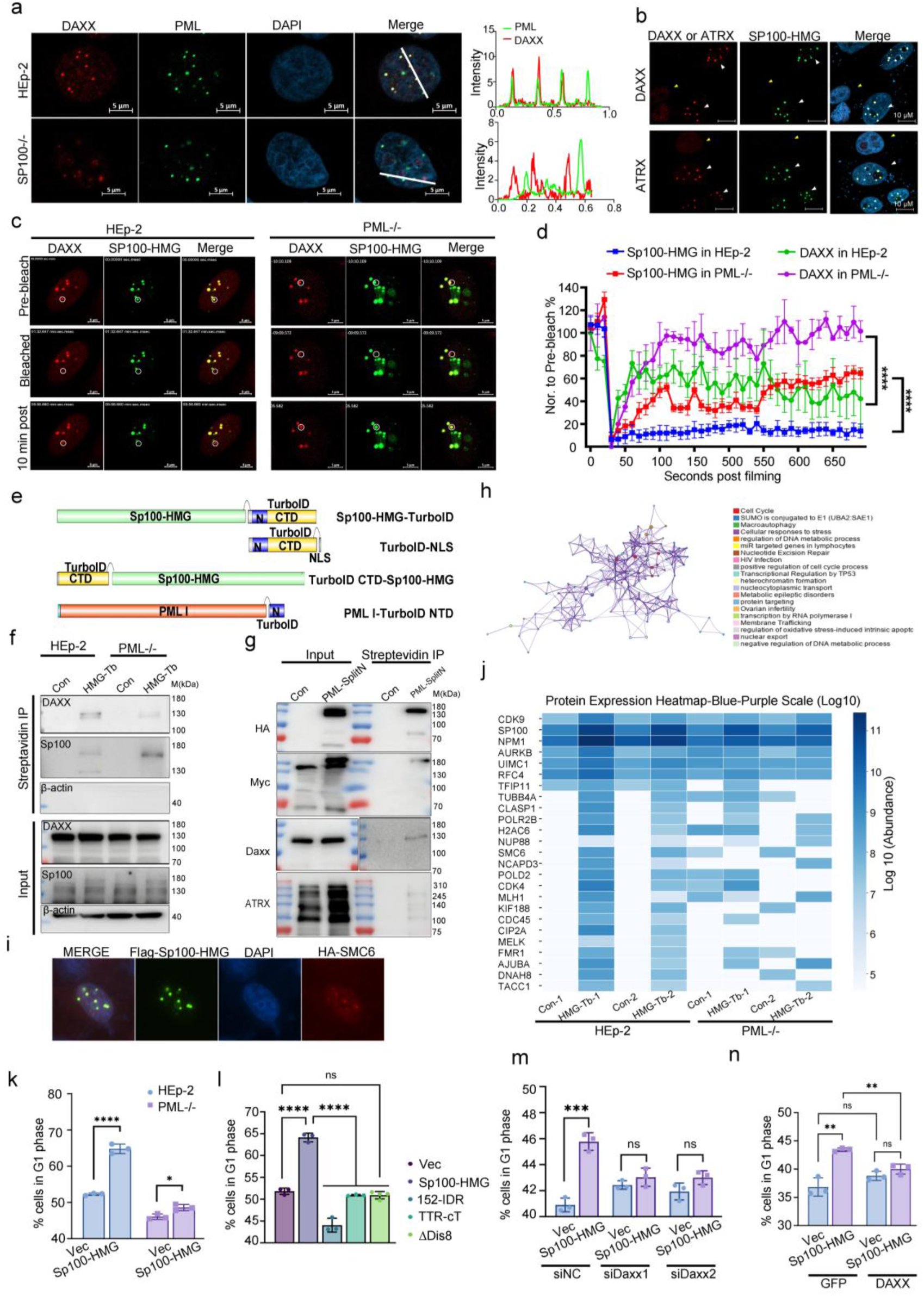
PML formed shell structure stabilize and enhanced client-protein recruitment in HMG-induced PML bodies that regulate cell cycle process in HEp-2 cells. A. Immunofluorescence of endogenous PML and DAXX in HEp-2 and Sp100-/- cells with corresponding line-scan fluorescence intensity profiles in the right panel. B. DAXX and ATRX localization in DKO cells w/wo Flag-Sp100-HMG (White arrows: positive cells; yellow arrows: negative cells). C-D. mCherry-DAXX was co-transfected with GFP-Sp100-HMG in HEp-2 and PML-/- cells for 24 hr. C. Representative FRAP time-lapse images in HEp-2 and PML-/- cells. D. Recovery kinetics of Sp100-HMG and DAXX in HEp-2 and PML-/- cells. E. Schematics of Sp100-HMG-TurboID and Split-TurboID constructs. F-G. Immunoblots of biotinylated proteins enriched via streptavidin beads immunoprecipitation from Sp100-HMG-TurboID (HMG-Tb) system in HEp-2 and PML-/- cells (F) or PML-I/Sp100-HMG Split-ID system in DKO cells (G). TurboID-NLS (con) served as control in F and TurboID-NLS (con) served as control in G. H. Metascape clusters of 103 Sp100-HMG-labeled proteins shared by 2 independently repeated mass spec results from Sp100-HMG-TurboID in HEp-2 cells. Node size reflects gene count; colors represent distinct biological pathways. J. Heatmap (z-score) showing abundance of 25 cell-cycle-associated proteins identified by mass spectrometry. K-N. Flow cytometry measuring G1 populations in HEp-2 or PML-/- cells following: (K) transfection of Sp100-HMG transfection, (L) transfection of Sp100-HMG mutants, (M) DAXX siRNA knockdown, or (N) DAXX overexpression.

To systematically map the proteomic constituents of Sp100-HMG condensates, we employed proximity labeling (TurboID and Split-ID) coupled with LC-MS/MS. We optimized these fusions to ensure that TurboID or Split-ID tags on Sp100-HMG and PML-I did not interfere with their LLPS capability while enabling efficient biotinylation (Fig 4 E, Fig S 5 D, E). Consistent with its role as a central scaffold, TurboID-Sp100-HMG effectively biotinylated DAXX in HEp-2 cells. However, this labeling efficiency was markedly reduced in PML-/- cells (Fig 4 F). Similarly, Split-ID successfully captured both DAXX and ATRX, providing a high-confidence map of the internal proteome (Fig. 4 G). LC-MS/MS identified 103 proteins significantly enriched (≥4-fold) in HMG-TurboID samples. MetaScape analysis revealed that these proteins are predominantly involved in cell cycle regulation and SUMOylation pathways (Fig 4 H, J, Fig S 5 F-H). We validated the recruitment of several identified candidates, including SMC6, SUMO1, and SUMO2, to Sp100-HMG puncta via imaging (Fig 4 I, Fig S 5 K).

Notably, the enrichment of cell cycle-related proteins was substantially disrupted in the absence of a functional PML shell (Fig. 4 J). This indicates that while Sp100-HMG serves as the autonomous scaffold for client recruitment, the PML shell is essential to maintain the biochemical density and spatial confinement of the interactome. Consistent with this model, key regulators such as CDK4 and CDK9 were prominently biotinylated by TurboID-Sp100-HMG and their labeling efficiency was significantly attenuated in PML-/- cells (Fig. S 5 I–J).

The enrichment of cell cycle regulators within the Sp100-HMG interactome led us to investigate its functional impact on cell progression. Flow cytometry revealed that Sp100-HMG overexpression induces a robust G1 phase arrest in HEp-2 cells, an effect entirely abolished in PML-/- cells (Fig. 4 K). Mutational analysis further demonstrated that this regulation requires the coordinated integrity of the "inside-out" assembly mechanism. Specifically, mutants defective in LLPS (ΔDis8), client sequestration (152-IDR, which fails to recruit DAXX), or PML shell organization (TTR-cT) all failed to induce cell cycle arrest without affecting DAXX levels (Fig. 4 L, Fig. S 5 L).

These results highlight a "double requirement": the Sp100-HMG core and the PML shell to provide the biochemical density required for functional sequestration. To test the "sponge" hypothesis—where the condensate depletes essential regulators from the nucleoplasm—we manipulated DAXX levels. DAXX knockdown abrogated the Sp100-HMG-induced G1 arrest phenotype (Fig. 4 M, Fig. S 5 M), while DAXX overexpression effectively counteracted the growth inhibition (Fig. 4 N, Fig. S 5 N).

Collectively, these results establish that Sp100-HMG condensates function as a phase-separated sponge that triggers G1 arrest by sequestering and depleting the functional nucleoplasmic pool of DAXX and other cell-cycle associated regulators. This identifies DAXX as the primary effector bridging the biophysical "inside-out" assembly of the nuclear body to physiological cell cycle control.

### Overexpression of functional Sp100-HMG in cancer cells hinders tumor growth in vivo

To investigate the tumor-suppressive potential of Sp100-HMG condensates, we systematically examined its effects across multiple tumor cell lines. Ectopic expression of Sp100-HMG induced robust growth inhibition in 12 of 20 tested cell lines, with efficacy strongly correlating with the formation of microscopically detectable nuclear condensates (Fig. S 6 and 7 A). Detailed analysis in H1299, T98G, and U2OS cells demonstrated significant proliferation inhibition at all measured timepoints (Fig 5A, Fig. S 7 A, B). The physiological relevance of these observations was supported by reciprocal experiments showing that siRNA-mediated knockdown of endogenous Sp100-HMG enhanced proliferation in both H1299 (Fig 5 B, C) and T98G cells (Fig. S 7 C).

**Fig. 5.**
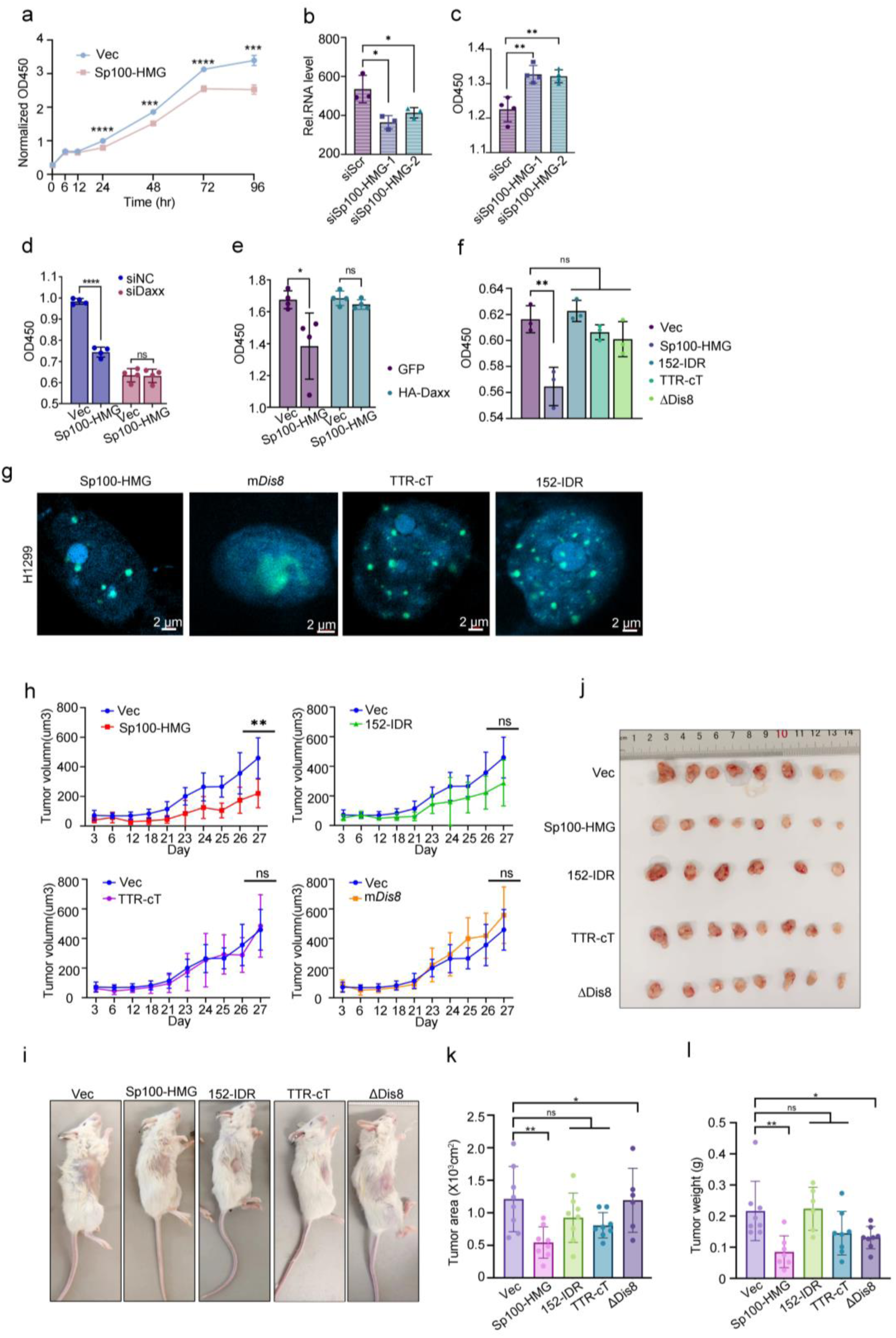
Sp100-HMG LLPS induced PML body organization suppressed tumor growth. A. CCK-8 proliferation curves of H1299 cells w/wo Sp100-HMG transfection. B-C. RSL3-induced endogenous Sp100-HMG expression followed by siRNA knockdown in H1299 showed inhibition of cell growth: qPCR validation (B) and CCK-8 proliferation at 36 hr (C). D-E. CCK-8 proliferation assays at 48 hr in H1299 cells, mirroring experimental conditions in Fig. 4 M-N. F-G. Proliferation rates of H1299 cells expressing various Sp100-HMG mutants (F) and corresponding immunofluorescence images of intranuclear patterns of Sp100-HMG and its mutants (G). H-I. Tumor growth curves and representative images of NCG mice subcutaneously injected with H1299 cells transfected with Sp100-HMG or mutants over 27 days. J-L. Images, volume measurements, and weights of excised tumors at day 27. The results were using unpaired two-tailed t-tests or one-way ANOVA where appropriate.

Importantly, the growth-inhibitory effect of Sp100-HMG overexpression in H1299 cells could be rescued by concurrent DAXX expression (Fig 5 D, E), mirroring our previous observations in HEp-2 cells (Fig 4 L). Functional dissection revealed that Sp100-HMG mutants defective in phase separation (ΔDis8), PML recruitment (TTR-cT), or DAXX binding (152-IDR) all failed to suppress proliferation in vitro (Fig 5 F, Fig. S 7 D).

These findings were further validated in an in vivo xenograft model using H1299 cells implanted in immunodeficient NCG mice (Fig 5 G). Tumors expressing wild-type Sp100-HMG showed significant reductions in both volume and weight compared to controls at all measured timepoints (Fig 5 H-L). In striking contrast, none of the functionally compromised Sp100-HMG variants exhibited tumor-suppressive activity, with xenografts showing comparable growth to vector controls. These results establish that Sp100-HMG-mediated tumor suppression requires the coordinated action of three essential molecular features: (1) liquid-liquid phase separation capacity conferred by its N-terminal multimerization domain and C-terminal disordered region, (2) SUMOylation-dependent integration into PML nuclear bodies, and (3) functional engagement of downstream effectors through its C-terminal interaction domain. The strict dependence on this tripartite mechanism across diverse experimental systems underscores the biological significance of Sp100-HMG condensate formation in growth control.

## Discussion

This study presents a ’core-nucleates-shell’ mechanism for de novo formation of PML NBs, suggesting that resident proteins, exemplified by Sp100-HMG, function as active drivers of architecture to initiate a multi-step phase separation process to organize these MLOs. These insights modify the decades-old dogma that PML is the sole organizer of the hierarchical assembly and illustrate how their distinct architecture underpins their functional versatility in transcriptional regulation, cell cycle control, and tumor suppression (93–95).

### Re-evaluating PML Nuclear Body Biogenesis

The canonical shell-first model faces fundamental biophysical paradoxes as previously discussed: the rigidity of the RBCC scaffold contradicts the compositional fluidity and rapid turnover of NBs, while the lack of autonomous PML-LLPS leaves the driving force for nucleation unexplained. These contradictions, combined with the observation of PML-independent cores (e.g., p62, Sp100), necessitate a paradigm shift (37, 62, 63). Rather than an autonomous scaffold, our findings extend this "inside-out" organizational model: an internal LLPS-driven condensate acts as the nucleation core, while the PML shell functions as a specialized interface or a surfactant-layer between the condensate and the nucleoplasm, thereby reconciling the stable structural shell with the dynamic internal environment. In this revised paradigm, the physical properties and identity of the internal protein condensate dictate the structural organization and functional output of the resulting mature MLO.

### Sp100-HMG as an Active Organizer of PML NB Assembly

Herein, we demonstrate that Sp100-HMG autonomously drives PML-NB formation via its intrinsic LLPS property. Domain analysis reveals that this process depends on the coordinated contributions of its N-terminal multimerization domain and C-terminal IDRs, where valency acts as the primary determinant over specific sequence for nucleation. Notably, nuclear membrane tethering assays underscore Sp100-HMG’s superior capacity to recruit PML compared to the inverse, positioning it as a critical nucleation factor.

We thus propose a novel ’core-nucleates-shell’ assembly model characterized by three hierarchical stages and redefines the role of the PML RBCC domain—traditionally viewed as the primary organizational driver—as a specialized factor for shell maturation rather than initial nucleation.

This modular framework intrinsically validates hallmark NB characteristics, such as internal fluidity and a concentric mobility gradient where the Sp100-HMG core restricts dynamics while client proteins like DAXX remain highly mobile. Furthermore, our discovery that ZBTB16 employs analogous mechanisms suggests this is a generalizable paradigm for PML-NB biogenesis. In this competitive and/or combinational nucleation model (Sp100-HMG, ZBTB16, and p62 serve as nucleating proteins), the local microenvironment dictates which component first reaches its critical saturation point (*Csat*); for instance, while p62 may template shells under stress, Sp100-HMG provides a lower kinetic barrier for nucleation in physiological or oncogenic contexts (63). Ultimately, this organization allows the PML shell to function as a flexible regulatory interface, while the specific internal core dictates the functional identity of the body across diverse nuclear landscapes.

### Functional Coupling of Sp100-HMG-PML NB Architecture to Regulation

Our findings demonstrate that the hierarchical "core-shell" architecture of PML NBs is intrinsically coupled to their regulatory output. We first show that artificially designed phase-separated condensates can template the recruitment of PML to specific genomic loci, functioning as transcriptional hubs and enhancing transcriptional activity of the transcriptional activator-rtTet, in a PML-dependent manner. Genome-wide mapping reveals that Sp100-HMG preferentially associates with transcriptional start sites, exerting context-dependent activation or repression. This spatial concentration of factors likely amplifies regulatory signals through the local enrichment of transcriptional machinery (Fig. 3K).

This architectural coupling extends to cell-cycle control. Proximity labeling identifies Sp100-HMG condensates as selective hubs for cell-cycle regulators, including DAXX, SMC6, and SUMO1/2. Crucially, the functional sequestering of these clients—leading to G1 arrest—requires a tripartite mechanism: Sp100-HMG-driven phase separation, proper PML recruitment, and functional client interaction. Disruption of any single component abolishes this regulatory effect, suggesting that the PML shell acts as a kinetic barrier that stabilizes the internal LLPS environment. This "protein sponge" effect allows for a tunable regulatory system, where the "breathing" behavior of the Sp100-HMG core—characterized by rhythmic expansion and contraction—modulates functioning site and nucleoplasmic protein availability in response to cellular cues, working in concert with the PML shell.

### Dynamic Heterogeneity and Scale-Dependent Functional Partitioning

The heterogeneity in PML NB size suggests a functional partitioning across different spatial scales. While research often focuses on micron-sized "visible" NBs, our framework hints at a vast population of sub-microscopic, "invisible" foci that likely execute localized, rapid functions such as DNA damage response, transcriptional control or ISG activation adjacent to chromatin (3, 96, 97). This is consistent with observations that core chromatin events, such as HIRA-mediated H3.3 deposition, can occur independently of large, juxtaposed NBs, pointing to the action of minimal, PML-dependent units (34).

In contrast, large visible NBs primarily function as "reservoirs" or "staging grounds" for protein sequestration, modification, or degradation. Our finding that PML-shelled Sp100-HMG condensates exert a "sponge effect" on DAXX aligns with the reported sequestration of HIRA for histone marking or p62 during cellular senescence (61). We propose that the biogenesis of PML NBs results from the collective participation of multiple LLPS-capable proteins, where local concentration thresholds dictate the transition from transient foci to robust, micron-sized regulatory hubs. While our reductionist approach using Sp100-HMG overexpression uncovers fundamental assembly rules, future investigations are needed to determine the interactive hierarchy connecting these diverse NB size classes across the nuclear landscape.

Collectively, our study employs a reductionist strategy—using tools like Sp100-HMG overexpression—to strip away cellular noise and uncover the fundamental mechanisms governing PML-NB assembly. These findings challenge the "shell-first" dogma and support a more dynamic, surfactant-mediated organizational principle. While we acknowledge the complexity of the native protein landscape, the biophysical constraints identified here serve as a critical foundation. They offer a new lens through which to investigate how the cell imposes spatial order upon the variable PML landscape, ultimately bridging the gap between molecular heterogeneity and architectural consistency.**Materials and Methods**

### Cell line and cultures

All cell lines used in this study were maintained in a humidified incubator at 37°C with 5% CO₂. Culture media were supplemented with 10% fetal bovine serum (FBS) unless otherwise specified. Cell lines were authenticated and cultured following ATCC or DSMZ guidelines and peer-reviewed protocols. The selection of basal media was based on standard requirements for each cell type, with tissue origin and culture background annotated as follows: Cells maintained in DMEM (High Glucose) included:HEp-2 (human epithelial carcinoma, historically associated with laryngeal origin), Beas-2B (normal human bronchial epithelium), MCF-7 and MDA-MB-231 (human breast adenocarcinoma), U87 (human glioblastoma), PC3 (human prostate adenocarcinoma), HELA (human cervical adenocarcinoma), B16 (murine melanoma), SW480 (human colorectal adenocarcinoma), and SW620 (human colorectal adenocarcinoma, lymph node metastasis). Additional cell lines including T98G, U251, LN229, DBTRG and M059K were also maintained in DMEM. Cells cultured in RPMI-1640 included: HCT116 (human colorectal carcinoma), H1299 (human non-small cell lung carcinoma), HEC-1A (human endometrial adenocarcinoma), and SW038 (assumed to refer to SW48, human colorectal adenocarcinoma). U2OS (human osteosarcoma) cells were cultured in McCoy’s 5A medium. Non-cancerous cell lines ARPE-19, HaCaT, and MRC-5 cell lines were cultured in DMEM. The glioma-related cell lines (U87, PC3, T98G, U251, LN229, DBTRG, M059K) and HCT116 were generously provided by Professors Deyin Guo and Xingan Fan (Sun Yat-sen University). ARPE-19 and NT2 cell line were obtained from Procell and AnWei Biotechnology Co., Ltd. HaCaT, and MRC-5 cell lines were kindly provided Doctor Bihui, Huang and Professor Hua, Zhang (Sun Yat-sen University). HEC-1A and MDA-MB-231 cells were obtained from Guangzhou Yuanjing Biotechnology Co., Ltd. Sp100 knockout (Sp100-/-), PML knockout (PM-/-), and double-knockout cell lines were generated using CRISPR/Cas9 genome editing as previously described (16, 17). For interferon stimulation, 24 hours post-transfection, the culture medium was replaced with fresh medium containing or lacking IFNβ (1000 U/mL) for an additional 24 hours. All cell lines were routinely tested for mycoplasma contamination.

### Plasmids and Transfection

All expression constructs used in this study were generated by subcloning the coding sequences into the pcDNA3.1 vector backbone. For fusion proteins, individual domains or tags were connected via a flexible protein linker. Site-directed mutants were generated using overlap extension PCR and confirmed by Sanger sequencing. Plasmids encoding GFP-RNF4 and GFP-UBC9 were purchased from Miaoling Biotechnology Co., Ltd.

Plasmids and siRNA were transfected into HEp-2 and other cell lines using JetPRIME reagent (Polyplus, Cat# 101000046) according to the manufacturer’s instructions. For plasmid transfection, cells were incubated for 24 hours before downstream assays. For siRNA-mediated knockdown experiments, cells were transfected and incubated for 36 hours unless otherwise specified. For NT-2 cell transfection: NT-2 cells (1 × 10⁶) were transfected with 1 µg of plasmid DNA using the CytoNanoTrans™ Transfection Reagent 3000.

### Tet-On expression system

The tet protein and the tetracycline-responsive promoter element (TRE) were cloned from the Tet-On expression system. Tet refers to the transactivator rtTA which contains a four amino acid change in the tetR DNA binding moiety that it can only recognize the tetO sequences in the TRE of the target transgene in the presence of the Dox effector

### Immunofluorescence Staining

Cells seeded on glass slides were washed three times with phosphate-buffered saline (PBS), then fixed with either ice-cold methanol (-20°C, 10 min) or 4% paraformaldehyde (PFA; RT, 15 min). After permeabilization with 0.3% Triton X-100 in PBS (RT, 10 min), cells were blocked with PBS-TBH blocking buffer (1× PBS containing 0.1% Tween-20, 10% FBS, and 3% bovine serum albumin) at room temperature for 30 min. Primary antibodies diluted in PBS-TBH were incubated overnight at 4°C or for 1 hr at 37°C. Following three PBS washes, samples were incubated with fluorophore-conjugated secondary antibodies Alexa Fluor 488 (Invitrogen, A32723) or Alexa Fluor 594 (Invitrogen, A11012) diluted in PBS-TBH for 30 min at 37°C in the dark. Slides were mounted with DAPI-containing medium (Abcam, ab104139) and imaged using a Zeiss LSM880 confocal microscope. High-resolution images were acquired using a Zeiss LSM 980 confocal microscope equipped with an Airyscan detector system.

### Live-Cell Condensate Dynamics and FRAP Analysis

To assess condensate dynamics in vivo, HEp-2 or other mammalian cell lines were transfected with GFP- or RFP-tagged constructs. After 24 hours, live-cell imaging was performed using a Zeiss LSM 880 confocal microscope. Fusion and fission events of nuclear or cytoplasmic puncta were monitored in real time. FRAP experiments were conducted under the appropriate laser excitation wavelengths by bleaching defined regions within fluorescent condensates. Fluorescence recovery was recorded over time, and recovery kinetics were analyzed using ImageJ software with the FRAP Profiler plugin to quantify intensity restoration.

To further assess the material properties of intracellular condensates, 1,6-hexanediol (1,6-HD) was added directly to the culture medium at a final concentration of 5% (v/v), and changes in condensate morphology and integrity were monitored by live imaging.

### TurboID-Based Proximity Labeling and Mass Spectrometry

To identify nuclear proteins in the vicinity of Sp100-HMG, we performed TurboID and Split-TurboID mediated proximity labeling, adapting the protocol established by Cho et al (98). For TurboID, cells were transfected with TurboID-Sp100-HMG fusion construct. Twenty-four hours after transfection, cells were incubated with 50 µM biotin for 10 minutes at 37°C to facilitate rapid biotinylation of proximal nuclear proteins. For SplitID, cells were co-transfected with TurboID CTD-Sp100-HMG and PML I-TurboID NTD (or control plasmid NLS-TurboID NTD) for 12 or 24 hr, then treated with 200 μM biotin for 15 min at 37°C. After labeling, cells were rinsed with cold PBS to halt the reaction and lysed in 1% SDS lysis buffer (50 mM Tris-HCl pH 7.5, 150 mM NaCl, 1 mM DTT, protease inhibitor cocktail). Biotinylated proteins were enriched by incubation with streptavidin-conjugated magnetic beads overnight at 4°C. Beads were washed stringently using buffers with high salt, alkaline pH, and urea to minimize nonspecific binding. Bound proteins were eluted in SDS loading buffer supplemented with free biotin and DTT, boiled for 10 minutes, and resolved by SDS-PAGE. Specific enrichment was confirmed by silver staining and streptavidin-HRP blotting. Gel slices corresponding to enriched bands in TurboID-Sp100-HMG samples were excised and submitted for LC-MS/MS analysis at the National Protein Science Facility, Tsinghua University. Data processing and protein identification were carried out using Pfind software.

### Metascape Analysis of the Mass Spectrometry data

Mass spec resulted protein abundances were compared as described in the corresponding result section. 103 proteins that were more than 4-fold enriched in HEp-2 cells transfected with Sp100-HMG-TurboID than those with NLS-TurboID were identified and analyzed in Metascape website (https://metascape.org). As according to the Metascape analysis process, they first identified all statistically enriched terms (can be GO/KEGG terms, canonical pathways, hall mark gene sets, etc., based on the default choices under Express Analysis or your choice during Custom Analysis) from the full cluster, accumulative hypergeometric p-values and enrichment factors were calculated and used for filtering. Remaining significant terms were then hierarchically clustered into a tree based on Kappa-statistical similarities among their gene memberships (similar to what is used in NCI DAVID site). Then 0.3 kappa score was applied as the threshold to cast the tree into term clusters. They selected a subset of representative terms from the full cluster and converted them into a network layout. More specifically, each term is represented by a circle node, where its size is proportional to the number of input genes fall under that term, and its color represent its cluster identity (i.e., nodes of the same color belong to the same cluster). Terms with a similarity score > 0.3 are linked by an edge (the thickness of the edge represents the similarity score). The network is visualized with Cytoscape with “force-directed” layout and with edge bundled for clarity. One term from each cluster is selected to have its term description shown as label (Fig 4 H). The same enrichment network has its nodes colored by p-value, as shown in the legend. The dark the color, the more statistically significant the node is (see legend for p-value ranges) (Sup Fig 5 G, H).

### Protein purification and in-vitro LLPS analysis of Sp100-HMG

The coding sequence of human Sp100-HMG was cloned into the pET-28a vector to generate an N-terminal His-tagged fusion protein. The construct was transformed into E. coli BL21 (AI) cells for expression. A single colony was cultured in LB medium at 37°C until OD₆₀₀ reached 0.5–0.6, followed by induction with 0.3 mM IPTG and overnight incubation at 18°C with shaking.

Cells were harvested by centrifugation (5,000 rpm, 15 min, 4°C) and resuspended in binding buffer (50 mM Tris-HCl pH 7.9, 500 mM NaCl, 10 mM imidazole). After lysis using a high-pressure homogenizer, the lysate was centrifuged at 18,000 rpm for 30 min at 4°C. The supernatant was incubated with Ni–NTA agarose beads for 30 min at 4°C, washed with binding buffer, and eluted with 500 mM imidazole. Proteins were further purified via size-exclusion chromatography using a Superdex 200 Increase column pre-equilibrated in high-salt (HS) buffer (25 mM HEPES pH 7.5, 500 mM NaCl, 1 mM DTT). Purified proteins were concentrated, aliquoted, flash-frozen, and stored at –80°C.

In vitro LLPS was induced by reducing ionic strength, increasing protein concentration, or adding PEG 8000 as a molecular crowding agent, following previously described protocols (67). Specifically, proteins were diluted from HS buffer to LLPS buffer (25 mM HEPES pH 7.5, 150 mM NaCl, 1 mM DTT) or supplemented with 5–20% (w/v) PEG 8000. Droplet formation was monitored by differential interference contrast (DIC) microscopy immediately after preparation. For biophysical assessment, fusion, wetting, and aging behaviors were examined. Sedimentation and salt-resistance assays were performed to determine solubility and phase transition properties. For fluorescence labeling, purified His-Sp100-HMG protein was labeled using the ALEXA Fluor 488 Microscale Protein Labeling Kit (Thermo Fisher Scientific, A30006) according to the manufacturer’s instructions. Labeled proteins were used for live imaging of phase-separated droplets.

FRAP (fluorescence recovery after photobleaching) analysis was performed using a Zeiss LSM 880 confocal microscope. Defined regions within protein droplets were bleached with high-intensity laser light, and fluorescence recovery was recorded over time to assess the dynamic properties of condensates. For disruption assays, 1,6-hexanediol (1,6-HD) was added to the droplet mixture at a final concentration of 5% (v/v). Droplet dissolution was observed in real time by DIC and fluorescence microscopy to evaluate the material state and reversibility of phase separation.

### CUT&Tag Assay

CUT&Tag experiments were carried out using the NovoNGS CUT&Tag 2.0 High-Sensitivity Kit (Novoprotein Scientific Inc, N259-YH01-01A), following the manufacturer’s protocol and prior reports (58). Briefly, 1 × 10⁵ HEp-2 cells or other cells, either unmodified or transfected with Flag-Sp100-HMG, were trypsinized and pelleted by centrifugation at 800 × g for 5 min at room temperature. Cells were bound to ConA-coated magnetic beads and suspended in 100 µL Dig-wash buffer (20 mM HEPES pH 7.5, 150 mM NaCl, 0.5 mM spermidine, protease inhibitors, and 0.05% digitonin) supplemented with 2 mM EDTA and primary anti-Flag antibody (1:100), followed by incubation at 4°C overnight. After triple washing with Dig-wash buffer, samples were incubated with a goat anti-mouse secondary antibody (Abcam, ab6708; 1:200) for 1 hr at room temperature. Following another three washes in Dig-HiSalt buffer (300 mM NaCl), protein A–Tn5 fusion enzyme was added and incubated at 37°C for 1 hr. Unbound enzyme was removed with Dig-HiSalt buffer. Tagmentation was triggered by adding 100 µL of tagmentation buffer (10 mM MgCl₂ in Dig-HiSalt buffer) and incubated at 37°C for 1 hr. Reactions were terminated with 0.5 M EDTA, 10% SDS, and proteinase K (20 mg/mL), followed by incubation at 55°C for 1 hr. DNA was purified using phenol–chloroform extraction and amplified via 15 cycles of PCR using barcoded primers. Libraries were cleaned using a 1.3:1 ratio of DNA clean beads to sample volume and quantified before pooling for sequencing.

### DNA Sequencing and Data Processing

Library size distribution and concentration were determined using the Qsep system. Paired-end sequencing was conducted on the Illumina platform according to the standard protocol. Raw reads were filtered with FASTp (v0.19.11) using the parameters: minimum length 50 and maximum number of N bases = 6, to remove adapters and low-quality sequences. Clean reads were aligned to the human reference genome (hg38) using BWA (v0.7.12-r1039) with options -T 25 -k 18. Only uniquely mapped reads were retained for downstream analysis. CUT&Tag signal profiles surrounding transcription start sites (±3 kb) were generated using the computeMatrix module in deepTools (v3.0.2). Peak calling was performed with MACS2 (v2.1.2) using the parameters: --qvalue 0.05 --call-summits --nomodel --shift -100 --extsize 200 --keep-dup all. Genomic annotation of peaks was carried out with ChIPseeker (69).

### RNA Extraction and Quantitative Real-Time PCR

Total RNA was isolated using either the Total RNA Extraction Kit (OMEGA, R6834) or TRIzol reagent, according to the respective manufacturer’s instructions. Complementary DNA (cDNA) synthesis was carried out with the Evo M-MLV Reverse Transcription Kit (Accurate Biology, AG11603). Quantitative real-time PCR (qRT-PCR) was performed using SYBR Green-based detection chemistry (Accurate Biology, AG11701) on a real-time PCR system. Primer sequences used for gene amplification are provided in Table 1.

### Cell Cycle and Cell Viability Analysis

HEp-2 or other mammalian cells were seeded into 6-well plates at approximately 70% confluence one day prior to transfection. After 12 hours, cells were transfected with the indicated plasmids and incubated for an additional 24 hours. For cell cycle analysis, transfected cells were harvested by trypsinization, washed twice with PBS, and fixed in 70% ethanol at 4°C overnight. Fixed cells were treated with RNase A (100 µg/mL) and stained with propidium iodide (PI, 50 µg/mL) in PBS for 30 minutes at room temperature in the dark. Flow cytometric analysis was performed using a BD FACSCanto II flow cytometer, and data were analyzed with FlowJo software to determine the percentage of cells in G0/G1, S, and G2/M phases.

For cell viability assays, transfected cells were counted and seeded into 96-well plates at a density of 10,000 cells per well. After 12 hours of attachment, 10 µL of CCK-8 reagent (Dojindo) was added to each well, followed by incubation for 2 hours at 37°C. Absorbance was measured at 450 nm using a BioTek microplate reader. Cell viability was normalized to control groups.

### Animal Experiments

Four- to six-week-old NCG mice were obtained from GemPharmatech (Nanjing, China). H1299 cells (2×10^6 per mouse), transfected with Sp100-HMG wild-type, its mutants, or control plasmids, were resuspended in 100 µL of sterile PBS and mixed with an equal volume of VITRO Gel at a 1:1 ratio. The mixture was incubated at room temperature for 15 minutes before subcutaneous injection into the flanks of NCG mice. Tumor growth was monitored over a 27-day period. The length and width of the tumors were measured externally at regular intervals using a digital caliper and the tumor volume was calculated using the formula: (π/6) × length × width². Tumor volume was plotted upon time using GraphPad Prism software. At the endpoint of the experiments, mice were euthanized, and tumor tissues were excised, weighed, photographed, and plotted. All experimental procedures were reviewed and approved by the Institutional Animal Care and Use Committee (IACUC) of Sun Yat-Sen University, under the approval number SYSU-IACUC-2024-003044.

### Statistical Analysis

All experiments were conducted with a minimum of three independent biological replicates unless otherwise stated. Data are expressed as mean ± standard deviation (SD). Statistical significance was assessed using two-tailed unpaired Student’s t-tests, or one-way/two-way ANOVA as indicated in figure legends, using GraphPad Prism 6.0. Significance was defined as follows: ns (not significant, P > 0.05), *P ≤ 0.05, **P ≤ 0.01, ***P ≤ 0.001, and ****P ≤ 0.0001.

If your research involved human or animal participants, please identify the institutional review board and/or licensing committee that approved the experiments. Please also include a brief description of your informed consent procure if your experiments involved human participants.

## Acknowledgments

This project was supported by the National Key Research and Development Program of China (2022YFC2305400), the National Natural Science Foundation of China (no. 31870157 and no. 32370161), the Shenzhen Science and Technology Innovation Program (JCYJ20180307151536743), Natural Science Foundation of Shenzhen City (JCYJ20220530145810023), Sanming Project of Medicine in Shenzhen (No. SZSM202411008), China Postdoctoral Science Foundation (2023M743996) and Guangdong Provincial Applied Science and Technology Research and Development Program (2025A1515012348).

## Author contributions

Conceptualization: P.X.

Methodology: P.X., H.D., Y.M., C.C.,

Investigation: H.D., Y.M., C.C., J.L., X.G., X.Z., W.L., X.D., L.Y., Y.L., H, L.

Visualization: P.X., H.D.

Funding acquisition: P.X. L.Y.

Project administration: P.X.

Supervision: P.X.

Writing – original draft: P.X., H.D.

Writing – review & editing: P.X., H.D.

## Data and materials availability

The data that support the findings of this study are available from the corresponding author, P.X., upon reasonable request.

