## Supporting text Figures S1 to S7 Table 1-3 for "Sp100-HMG drives ‘inside-out’ PML-NB assembly to modulate transcription and cell-cycle dynamics"

**This PDF file includes:**

Supporting text
Figures S1 to S7
Table 1-3

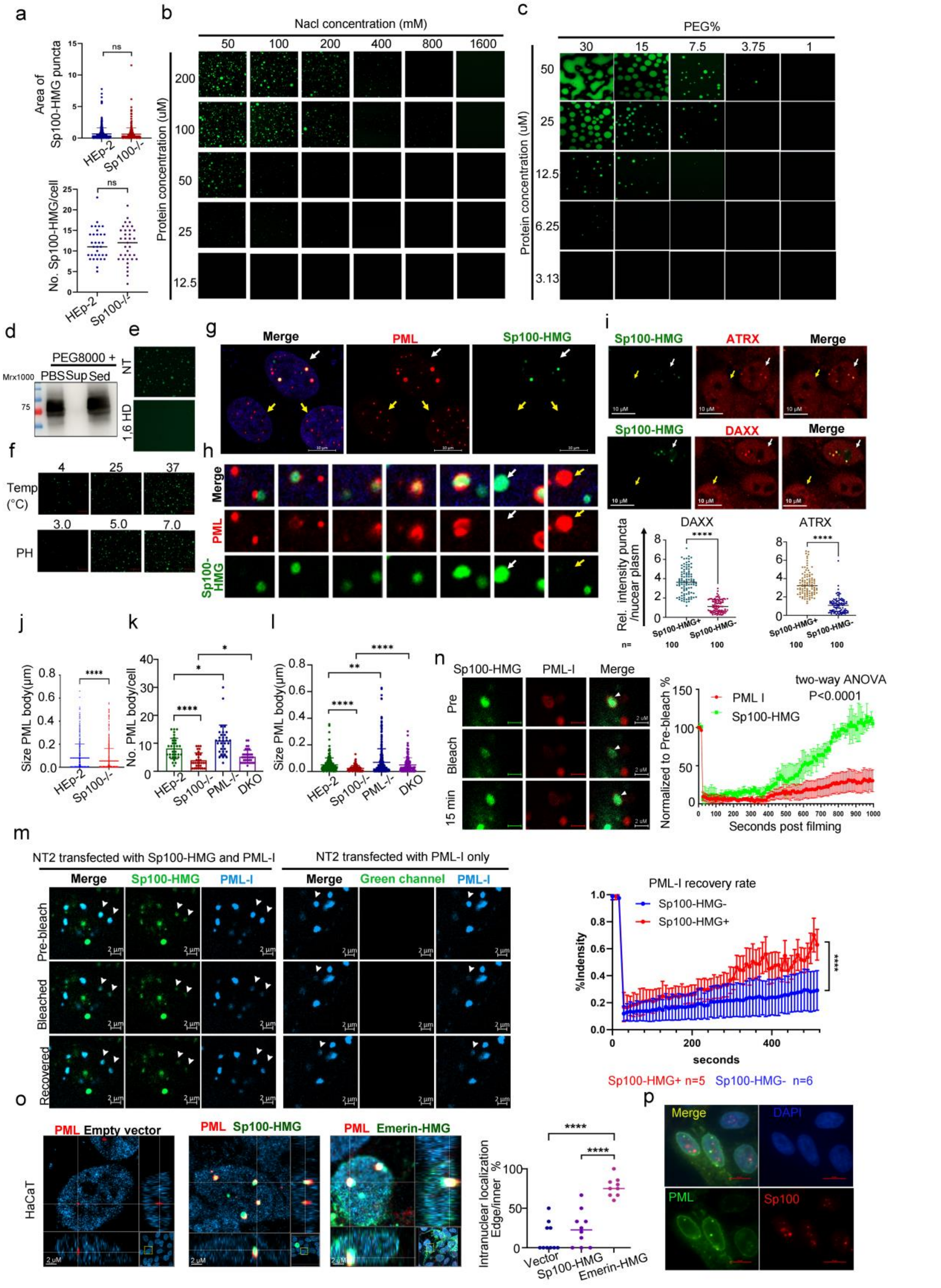


**Fig. S 1.** A. Characterization of the Sp100-HMG speckles in the Sp100-HMG transfected HEp-2 and Sp100-/- cells (Up: area, down: number). B-C. Bacterially purified His-Sp100-HMG protein was labeled using the ALEXA Fluor 488 Microscale Protein Labeling Kit. Representative images of purified Sp100-HMG at the indicated protein concentration and iron strength (B) or PEG concentration (C). D. Bacterially purified Sp100-HMG was dissolved in PBS or PBS supplemented with 15% PEG8000 and spined at 8000Xg for 10 min. Sp100-HMG levels in the PBS only or in the supernatant and sediment of PEG8000 contained solution were analyzed by immunoblot. E. Representative images showing the Sp100-HMG condensates treated with (1,6-HD) or without 1,6 HD (NT). F. Representative images showing formation of the Sp100-HMG condensates at indicated temperature and the indicated PH. G-I. Cells were transfected with Flag-tagged Sp100-HMG for 24 hr. NT2 cell were fixed and immune-stained for Sp100-HMG and PML (G). Representative images are shown presenting both Sp100-HMG positive cells (white arrows) and Sp100-HMG negative cells (yellow arrows). Different colocalization patterns of Sp100-HMG and PML were shown (H). NT2 cells were immune-stained for transfected Sp100-HMG, endogenous DAXX and ATRX. Quantification of the fluorescence intensities of DAXX and ATRX within the condensates versus the surrounding nucleoplasm for both Sp100-HMG positive and negative cells is presented in the lower panel. J-L. Quantification of the endogenous PML body diameters in HEp-2 and Sp100-/- cells (J). Quantification of the PML body number (K) and size (L) in the indicated cells transiently transfected with PML-I. M. NT2 cells were transfected with BFP- PML-I alone or together with GFP-Sp100-HMG for 24 hr. Representative time-lapse images showing FRAP results of BFP-PML-I at the indicated time points (bleached spots indicated by white arrow heads) (Left). Recovery kinetics of PML (Right). N. A defined region (white arrow head) within the condensate formed by GFP-tagged Sp100-HMG and mCherry tagged PML-I in DKO cells was photobleached, and fluorescence recovery was monitored over time. Representative time-lapse images at the indicated time points are shown in L and fluorescence recovery kinetics of PML-I and Sp100-HMG are plotted in right panel. O. HaCaT cells were transfected with siSp100 and then with empty vector, Sp100-HMG or Emerin-Sp100-HMG (Emerin-HMG) for 24 hr, fixed and immune-stained for endogenous PML and Sp100-HMG. Representative fluorescence images and graphic illustrations showing endogenous PML and transfected Sp100-HMG-Emerin or Sp100-HMG staining (Left). Percentage of nuclear edge located PML speckles was quantified and plotted (Right). P. DKO cells were transfected with Emerin-PML-I and Sp100-HMG for 24 hr, cells were fixed and immune-stained for PML and Sp100. Representative images showing the localization of Emerin-PML-I and Sp100-HMG.

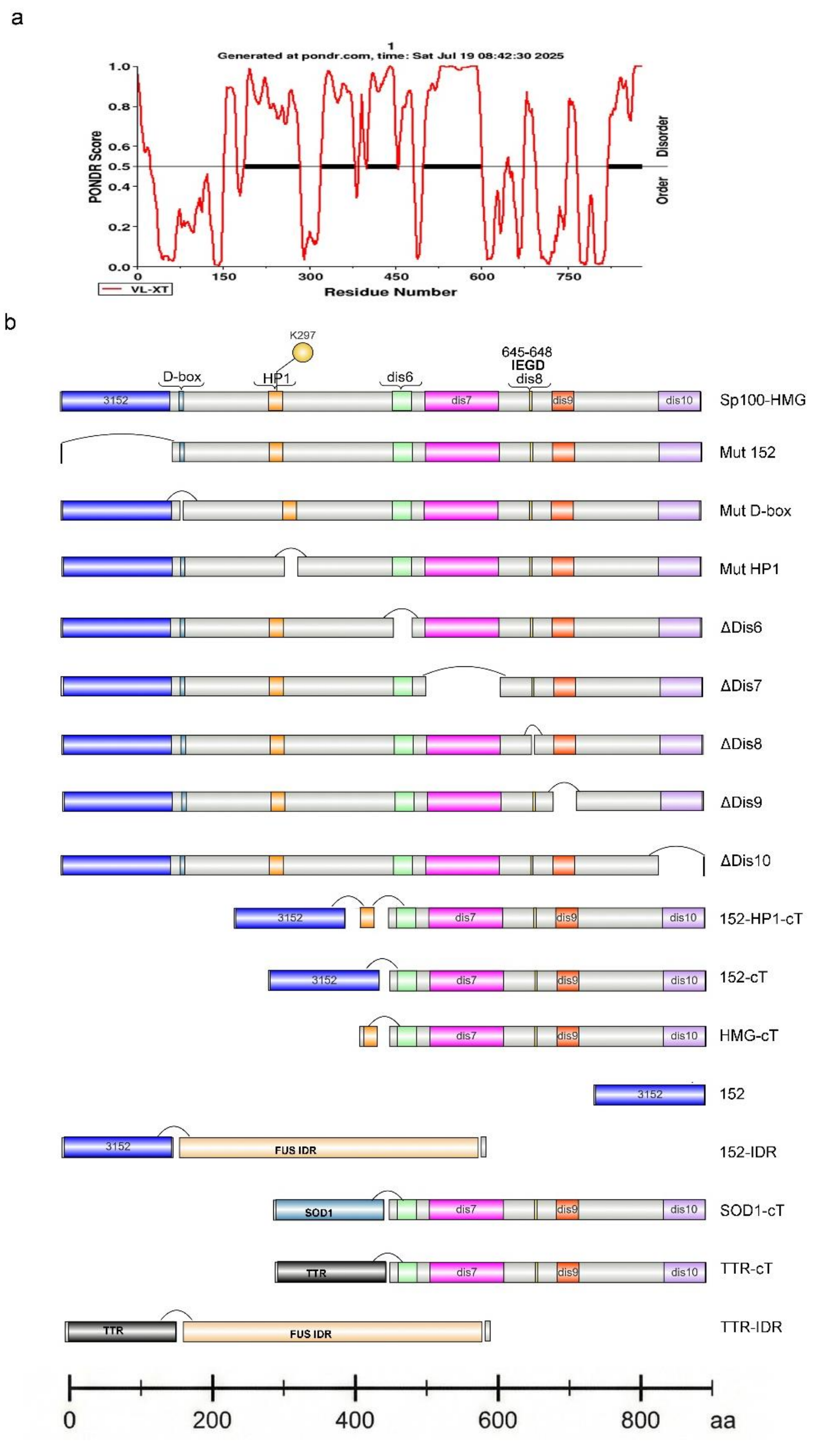


**Fig. S 2.** A. Predicted IDR regions within the C terminal of Sp100-HMG. B. Illustrations of Sp100-HMG, its variants and artificially constructed LLPS inducers used in this study.

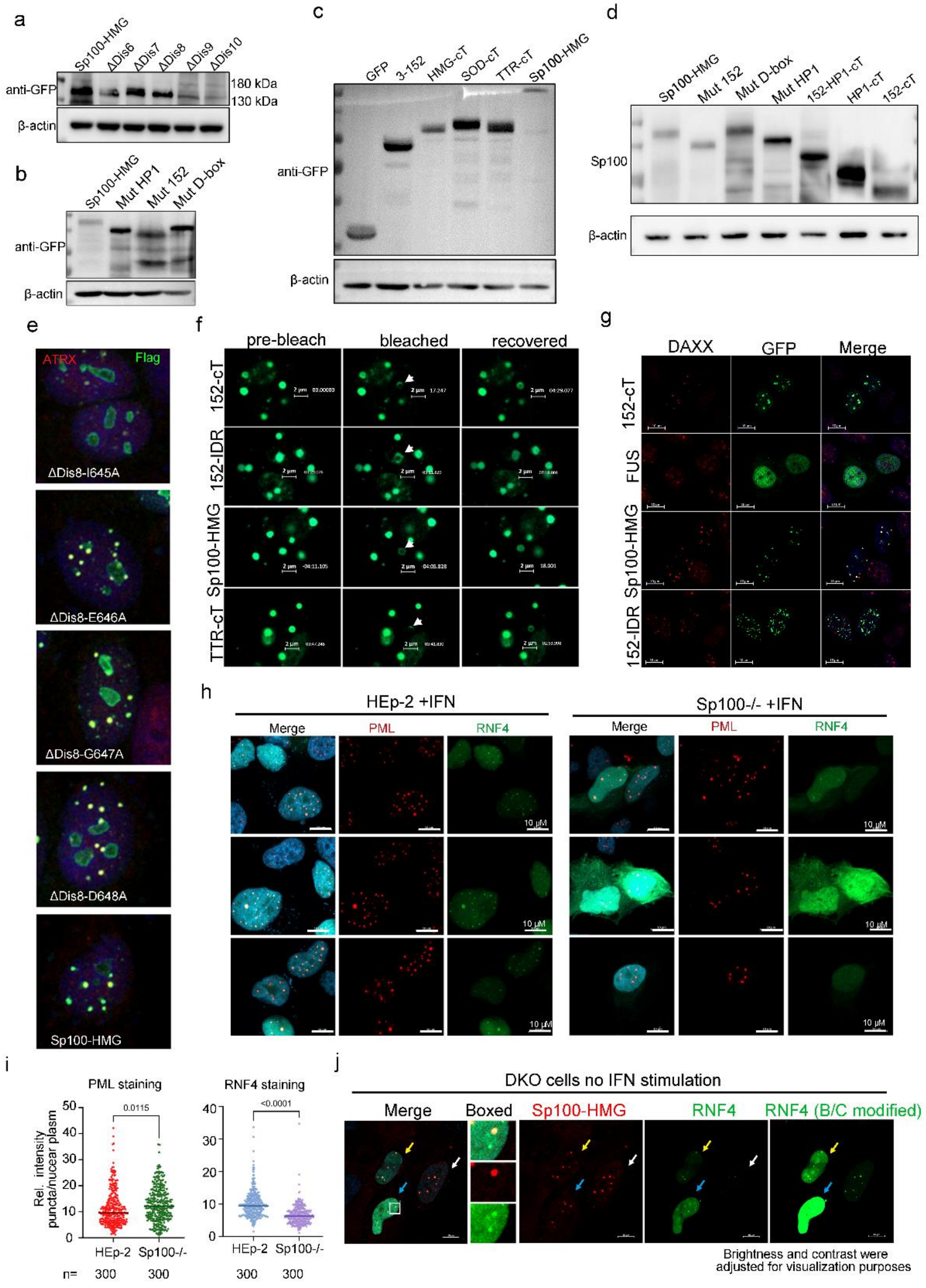
a
Sp100-HMG
ΔDis6
ΔDis7
ΔDis8
ΔDis9
ΔDis10
anti-GFP
180 kDa
130 kDa
β-actin
b
Sp100-HMG
Mut HP1
Mut 152
Mut D-box
anti-GFP
β-actin
c
GFP
3-152
HMG-cT
SOD-cT
TTR-cT
Sp100-HMG
anti-GFP
β-actin
d
Sp100-HMG
Mut 152
Mut D-box
Mut HP1
152-HP1-cT
HP1-cT
152-cT
Sp100
β-actin
e
ATRX
Flag
ΔDis8-I645A
ΔDis8-E646A
ΔDis8-G647A
ΔDis8-D648A
Sp100-HMG
f
pre-bleach
bleached
recovered
152-cT
152-IDR
Sp100-HMG
TTR-cT
2 μm
g
DAXX
GFP
Merge
152-cT
FUS
Sp100-HMG
152-IDR
h
HEp-2 +IFN
Sp100-/- +IFN
Merge
PML
RNF4
10 μM
i
PML staining
RNF4 staining
Rel. intensity puncta/nucear plasm
0.0115
<0.0001
HEp-2
Sp100-/-
n=
300
300
300
300
j
DKO cells no IFN stimulation
Merge
Boxed
Sp100-HMG
RNF4
RNF4 (B/C modified)
Brightness and contrast were adjusted for visualization purposes

k

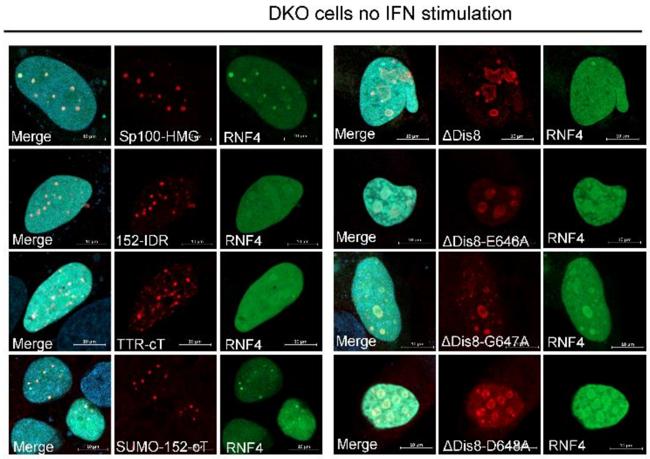


l

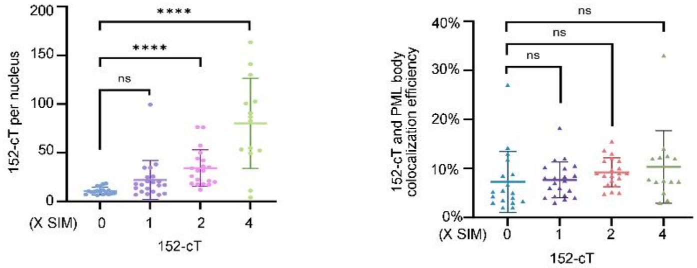


m

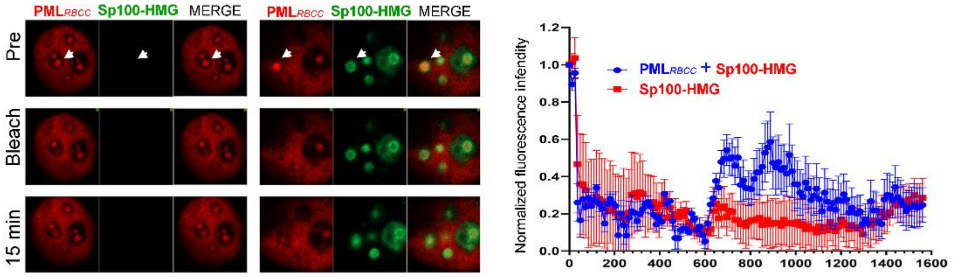


**Fig. S 3.** A-D. Western blot analysis of Sp100 mutants' expression in HEp-2 cells. A. Sp100-HMG and the IDR mutants including ΔDis6, ΔDis7, ΔDis8, ΔDis9, ΔDis10; B. Sp100-HMG, and the N terminal mutants including Mut 152, Mut HP1 and Mut D-box; C. GFP and fusion constructs 3-152, HMG-cT, SOD-cT, TTR-cT and Sp100-HMG; D. Sp100-HMG and fusion constructs Mut 152, Mut D-box, Mut HP1, 152-HP1-cT, HP1-cT and 152-cT. E. Individual amino acid mutations in the Dis8 region were constructed and transfected into DKO cells. After 24 hr transfection, cells were fixed and detected with IF using anti-Flag antibodies. F. Representative time-lapse images of FRAP on GFP-Sp100-HMG or its GFP tagged mutants in DKO cells. G. Evaluation of endogenous DAXX recruitment by the indicated mutants. H-I. HEp-2 and Sp100-/- cells transfected with GFP-RNF4 were treated with IFNβ for 24 hr, fixed and immune-stained for PML and RNF4. Representative images were shown in H. The fluorescence intensities of PML and RNF4 within the condensates versus the surrounding nucleoplasm was quantified and plotted in I. J-K. DKO cells were transfected with Sp100-HMG or the indicated Sp100-HMG mutants and GFP-RNF4, fixed and immune-stained with proper antibodies. J. White, yellow and blue arrows indicate cells with different expression levels of RNF4. L. Quantification of the number of puncta per nucleus for the 152-cT construct containing the indicated number of SIM repeats (Left). Quantification of colocalization efficiency of PML and the 152-cT construct containing the indicated number of SIM repeats (Right).

M. A defined region within the condensate formed by mCherry tagged $PML_{RBCC}$ with or without GFP-Sp100-HMG were photobleached, and fluorescence recovery was monitored over time. Left, representative time-lapse images of FRAP in DKO cells. Right, Recovery kinetics of $PML_{RBCC.}$

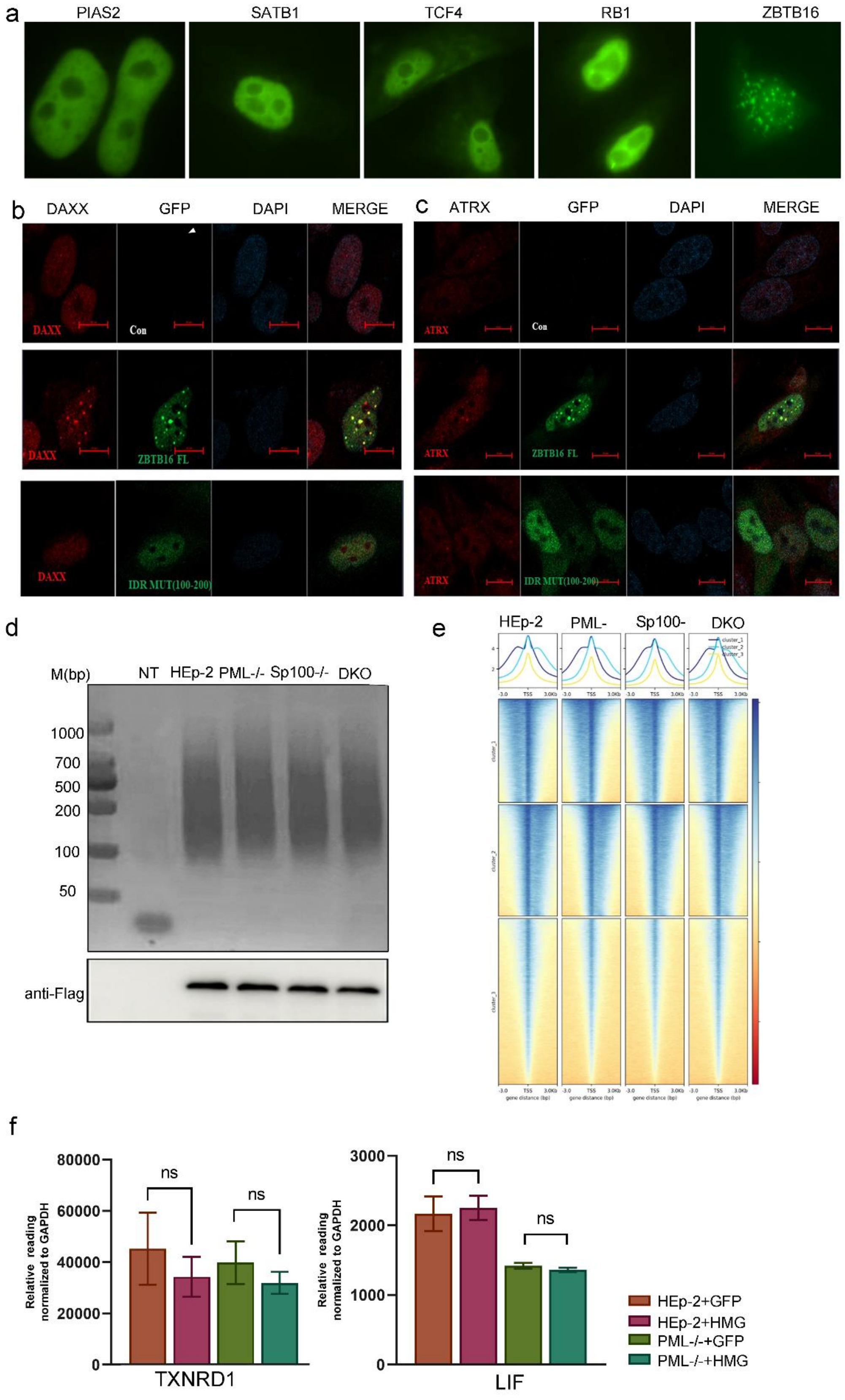
a
PIAS2
SATB1
TCF4
RB1
ZBTB16
b
DAXX
GFP
DAPI
MERGE
DAXX
Con
DAXX
ZBTB16 FL
DAXX
IDR MUT(100-200)
c
ATRX
GFP
DAPI
MERGE
ATRX
Con
ATRX
ZBTB16 FL
ATRX
IDR MUT(100-200)
d
M(bp)
NT
HEp-2
PML-/-
Sp100-/-
DKO
1000
700
500
200
100
50
anti-Flag
e
HEp-2
PML-
Sp100-
DKO
f
80000
60000
40000
20000
0
Relative reading normalized to GAPDH
ns
ns
TXNRD1
3000
2000
1000
0
ns
ns
LIF
HEp-2+GFP
HEp-2+HMG
PML-/-+GFP
PML-/-+HMG

**Fig. S 4.** A. The indicated proteins were cloned, tagged to GFP and transfected in DKO cells. Representative images are shown. B-C. Colocalization transfected ZBTB16 or its mutants with endogenous DAXX (B) and ATRX (C) in DKO cells. D. Assessment of the Cut&Tag libraries by agarose gel electrophoresis. Expression of the Sp100-HMG in the indicated cell lines was detected by immunoblot using anti-Flag antibody. E. Heat map showing the distribution of Sp100-HMG on genomes in the indicated cell lines. F. Sp100-HMG or GFP expressing plasmid was transfected into HEp-2 and PML-/- cells, RNA levels of the indicated genes were quantified by qRT-PCR and normalized to GAPDH.

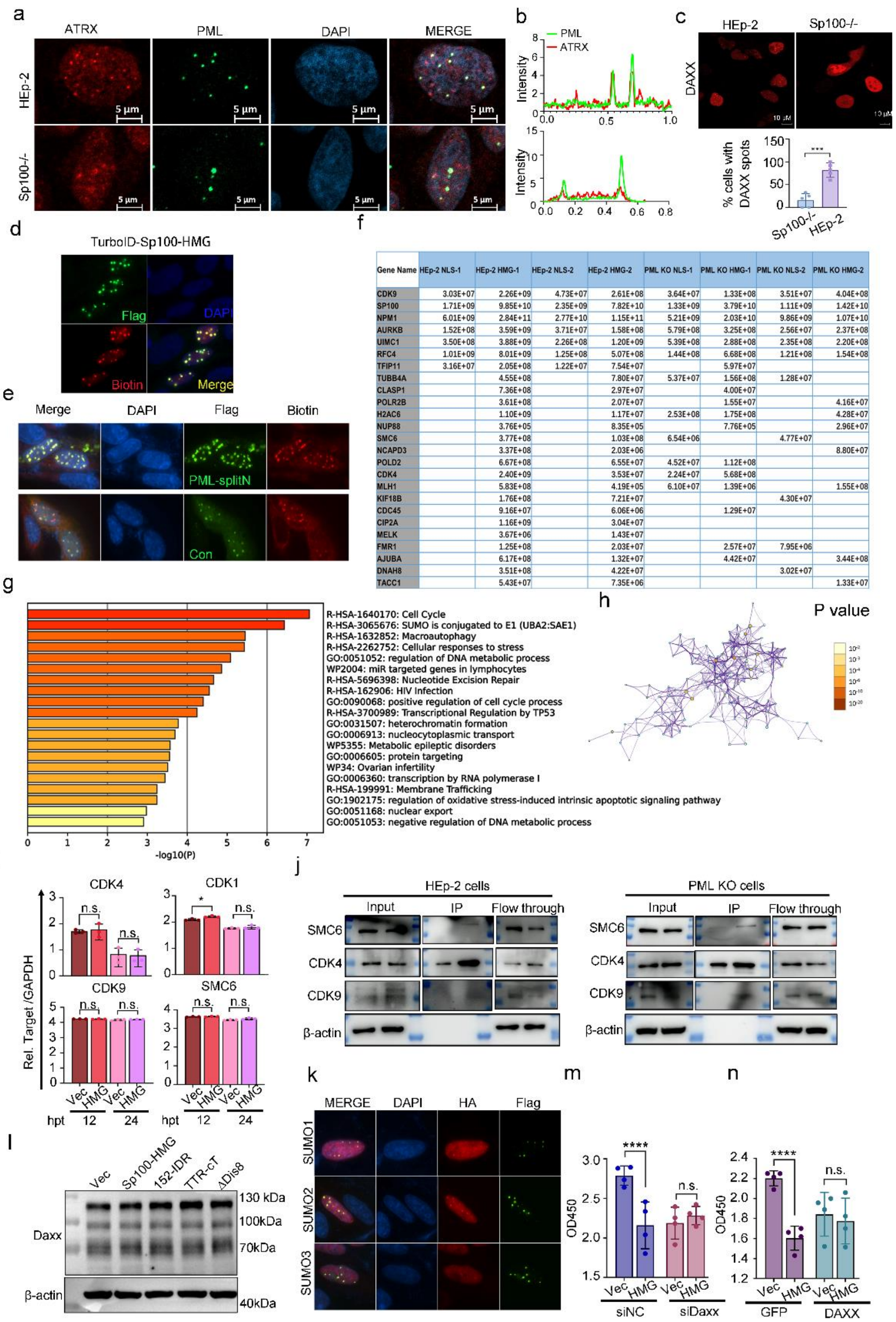


| Gene Name | HEp-2 NLS-1 | HEp-2 HMG-1 | HEp-2 NLS-2 | HEp-2 HMG-2 | PML KO NLS-1 | PML KO HMG-1 | PML KO NLS-2 | PML KO HMG-2 |
|---|---|---|---|---|---|---|---|---|
| CDK9 | 3.03E+07 | 2.26E+09 | 4.73E+07 | 2.61E+08 | 3.64E+07 | 1.33E+08 | 3.51E+07 | 4.04E+08 |
| SP100 | 1.71E+09 | 9.85E+10 | 2.35E+09 | 7.82E+10 | 1.33E+09 | 3.79E+10 | 1.11E+09 | 1.42E+10 |
| NPM1 | 6.01E+09 | 2.84E+11 | 2.77E+10 | 1.15E+11 | 5.21E+09 | 2.03E+10 | 9.86E+09 | 1.07E+10 |
| AURKB | 1.52E+08 | 3.59E+09 | 3.71E+07 | 1.58E+08 | 5.79E+08 | 3.25E+08 | 2.56E+07 | 2.37E+08 |
| UIMC1 | 3.50E+08 | 3.88E+09 | 2.26E+08 | 1.20E+09 | 5.39E+08 | 2.88E+08 | 2.35E+08 | 2.20E+08 |
| RFC4 | 1.01E+09 | 8.01E+09 | 1.25E+08 | 5.07E+08 | 1.44E+08 | 6.68E+08 | 1.21E+08 | 1.54E+08 |
| TFIP11 | 3.16E+07 | 2.05E+08 | 1.22E+07 | 7.54E+07 |  | 5.97E+07 |  |  |
| TUBB4A |  | 4.55E+08 |  | 7.80E+07 | 5.37E+07 | 1.56E+08 | 1.28E+07 |  |
| CLASP1 |  | 7.36E+08 |  | 2.97E+07 |  | 4.00E+07 |  |  |
| POLR2B |  | 3.61E+08 |  | 2.07E+07 |  | 1.55E+07 |  | 4.16E+07 |
| H2AC6 |  | 1.10E+09 |  | 1.17E+07 | 2.53E+08 | 1.75E+08 |  | 4.28E+07 |
| NUP88 |  | 3.76E+05 |  | 8.35E+05 |  | 7.76E+05 |  | 2.96E+07 |
| SMC6 |  | 3.77E+08 |  | 1.03E+08 | 6.54E+06 |  | 4.77E+07 |  |
| NCAPD3 |  | 3.37E+08 |  | 2.03E+06 |  |  |  | 8.80E+07 |
| POLD2 |  | 6.67E+08 |  | 6.55E+07 | 4.52E+07 | 1.12E+08 |  |  |
| CDK4 |  | 2.40E+09 |  | 3.53E+07 | 2.24E+07 | 5.68E+08 |  |  |
| MLH1 |  | 5.83E+08 |  | 4.19E+05 | 6.10E+07 | 1.39E+06 |  | 1.55E+08 |
| KIF18B |  | 1.76E+08 |  | 7.21E+07 |  |  | 4.30E+07 |  |
| CDC45 |  | 9.16E+07 |  | 6.06E+06 |  | 1.29E+07 |  |  |
| CIP2A |  | 1.16E+09 |  | 3.04E+07 |  |  |  |  |
| MELK |  | 3.67E+06 |  | 1.43E+07 |  |  |  |  |
| FMR1 |  | 1.25E+08 |  | 2.03E+07 |  | 2.57E+07 | 7.95E+06 |  |
| AJUBA |  | 6.17E+08 |  | 1.32E+07 |  | 4.42E+07 |  | 3.44E+08 |
| DNAH8 |  | 3.51E+08 |  | 4.22E+07 |  |  | 3.02E+07 |  |
| TACC1 |  | 5.43E+07 |  | 7.35E+06 |  |  |  | 1.33E+07 |

**Fig. S 5.** A-B. HEp-2 cells and Sp100-/- cells were staining for endogenous PML and ATRX with proper antibodies. A. Representative images. B. Fluorescence intensity along the white line in A were measured by image J and plotted. C. HEp-2 and Sp100-/- cells were transfected with a DAXX-expressing plasmid for 24 hr and analyzed for the intranuclear distribution pattern of DAXX. Representative images are shown, and the percentage of cells exhibiting puncta-associated DAXX is plotted in the lower panel. (n=54 in HEp-2, n=43 in Sp100-/-). D. HEp-2 cells transfected with

Flag-Sp100-HMG-TurboID were supplemented with biotin as in Figure 4 F, fixed and stained against biotin and Flag tag. E. PML-/- cells transfected with Flag-TurboID CTD-Sp100-HMG and NLS-TurboID NTD (con) or PML I- TurboID NTD (PML-SplitN) were supplemented with biotin as in Figure 4 F, fixed and stained with anti-streptavidin and -Flag antibodies. F. The relative peptide abundance readings of 25 cell cycle associated proteins intensively labeled by Sp100-HMG TurboID in HEp-2 cells are shown. G-H. As described in Figure 4 H, 103 genes were analyzed for their biological functions using Metascape, enriched ontology clusters across studies are shown in G and enriched ontology clusters colored by p-value is shown in H. I. HEp-2 cells were transfected with empty vector or Sp100-HMG for 12 hr and 24 hr, mRNA expression levels of the indicated genes were quantified by qRT-PCR and normalized to GAPDH. J. TurboID labeling by Sp100-HMG was conducted in HEp-2 and PML KO cells as in Figure 4 F. Biotinylated proteins in the cell lysate (Input) and during enrichment by streptavidin agarose beads (IP and Flowthrough) were immunoblotted with the indicated antibodies. K. Co-localization of HA-SUMO1, SUMO2 and SUMO3 with Flag-Sp100-HMG in HEp-2 cells by transfection and immunofluorescence labeling. L. HEp-2 cells were transfected with empty vector or Sp100-HMG for 24 hr, protein level of DAXX was confirmed by immunoblot. M-N. Experiments were conducted as in Figure 4 M and N, cell proliferation rates at 36 hr post transfection were measured by CCK-8 assay and absorbance at OD450 were plotted.

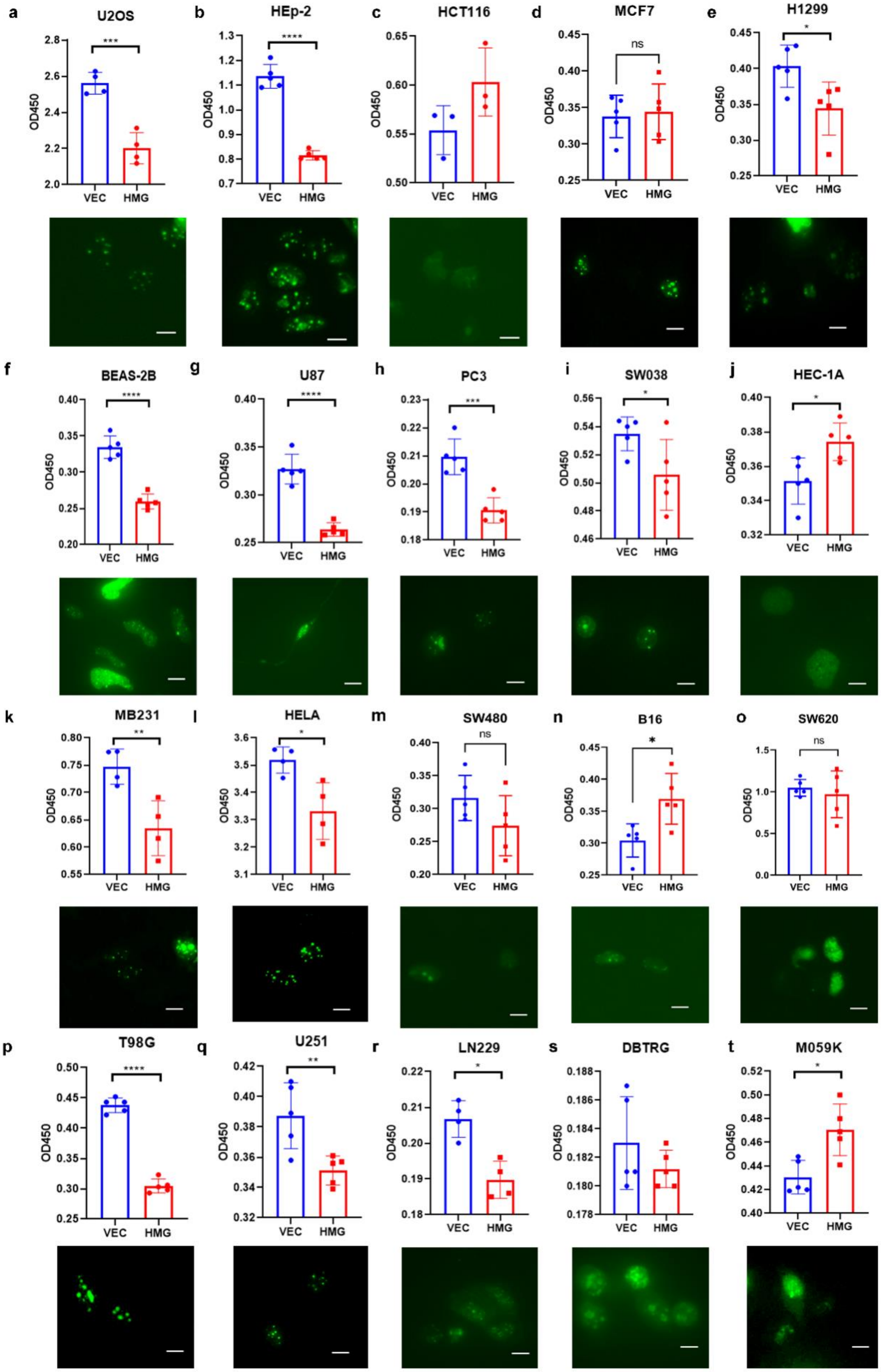
a
U2OS
***
OD450
VEC
HMG
b
HEp-2
****
c
HCT116
d
MCF7
ns
e
H1299
*
f
BEAS-2B
****
g
U87
****
h
PC3
***
i
SW038
*
j
HEC-1A
*
k
MB231
**
l
HELA
*
m
SW480
ns
n
B16
*
o
SW620
ns
p
T98G
****
q
U251
**
r
LN229
*
s
DBTRG
t
M059K
*

**Fig. S6.** A-T. Tumor cell lines were transfected with GFP-Sp100-HMG expression plasmid for 24 hr. The intranuclear pattern of GFP-Sp100-HMG was imaged and shown at the lower part in each figure panel and the impact of its overexpression on cell proliferation was measured by CCK8-assay and shown at the top part. Scale bar = 5 μM.

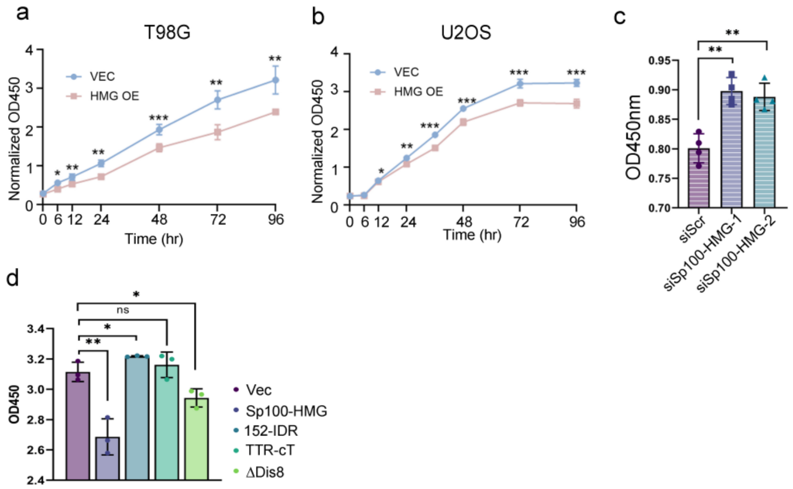


**Fig. S7.** A. T98G cells were transfected with Sp100-HMG or control vector and cell proliferation level at the indicated time points was measured by CCK-8 assay and absorbance at OD450 were plotted. B. Same experiments were conducted in U2OS cells. C.T98G cells were treated with RSL3 to enhance transcription from Sp100 gene and transfected with scramble siRNA (siRNA) or siRNAs specifically targeting Sp100-HMG isoforms (siSp100-HMG-1 and -2) and cell proliferation rate was measure by CCK-8 assay at 48 hr post transfection. D. T98G cells were transfected with Sp100-HMG or the indicated mutant constructs. Cell proliferation rate was assessed by CCK-8 assay.

**Tables**

Table 1. Primers for qRT-PCR.

| Gene | Forward | Reverse |
|---|---|---|
| PIEZO1 | AGGCGCATCAGTCTACGTTT | GCTTGGCCTCTTCTCTCTCC |
| NOD1 | TACTGAAAAGCAATCGGGAACT | GTAGAGGAAGAACTCGGACACC |
| EPCAM | AATCGTCAATGCCAGTGTACTT | TCTCATCGCAGTCAGGATCATAA |
| LIF | CGAGCCTATACAGATGGTGGA | CCATTCTCGTTTCCGATA |
| TXNRD1 | ATGGCAAGAAGGTGATGG | GCAGTAAGGCAAGGAGAA |
| EGFR | AGGCACGAGTAACAAGCTCAC | ATGAGGACATAACCAGCCACC |
| Firefly Luciferase | ATAACTGGTCCGCAGTGGTG | AGGCCGCGTTACCATGTAAA |
| Rellina Luciferase | GAGGCGAACTGTGTGTGAGA | GTGTTCGTCTTCGTCCCAGT |
| Sp100-HMG | AGCTAAGTATACGCTGCGGT | ACTTGACTGAAGCATCTGGGT |
| CDK1 | GGCTCTGATTGGCTGCTTTG | GGTAGATCCGCGCTAAAGGG |
| CDK4 | ATGGCTGCCACTCGATATGAACCC | GTACCAGAGCGTAACCACCACAGG |
| CDK9 | AAAACGAGAAGGAGGGGTTCC | CCTTGCAGCGGTTATAGGGG |
| SMC6 | CCTAAAAATGCCAAAAGGCCAAG | TTTACATTCGTCTTCGTCACCAT |
| GAPDH | ATGACATCAAGAAGGTGGTG | CATACCAGGAAATGAGCTTG |

Table 2. Primers for PCR.

| Gene | Forward | Reverse |
|---|---|---|
| SP100-HMG | TTAGGCGATATCCCGTTATTTATCATCATCTTCTTCA | GCGTTAGATATCCGCTCACTTGATCATCACCTT |
| DIS6 | GGTGGAGGTGGTTCTGGTGGAGGTGGATCTCCTGAAAATAAGAAGTGCTC | CCACCAGAACCACCTCCACCTGAAAAGTCACTACTGCTGAAA |
| DIS7 | GTTCTGGTGGAGGTGGATCTCAATCTGAACTTCCTGTGAC | AGATCCACCTCCACCAGAACCACCTCCACCTGGCACACCTTTTGGAAAAC |
| DIS8 | AATTTGAAGCAGCAGCAGCACGCGGAGCATCCAAGAACTG | TGCTGCTGCTGCTTCAAATTCCCTGGGAGT |
| DIS9 | GGTGGAGGTGGTTCTGGTGGAGGTGGATCTGTTGACCCTTGTGAGGAGCA | CCACCAGAACCACCTCCACCAAATTTGTTCTCCATCAGGA |

| | | |
|---|---|---|
| DIS10 | TTAGGCGATATCCCGTTATTTATCATCATCTTCTTCA | ACAAGCAGTTTTATGAAAAGTAACGGGATATCCAGC |
| D8-1 | GAATTTGAAGCAGAAGGAGACCGC | TCCGCGGTCTCCTTCTGCTTCAAA |
| D8-2 | GAATTTGAAATTGCAGGAGACCGC | TCCGCGGTCTCCTGCAATTTCAAA |
| D8-3 | GAATTTGAAATTGAAGCAGACCGC | TCCGCGGTCTGCTTCAATTTCAAA |
| D8-4 | TTTGAAATTGAAGGAGCACGCGGA | TGCTCCGCGTGCTCCTTCAATTTC |
| TURBOID-NLS | AGCTGCTAGCGCCATGAAAGACAATACTGTGCCTCT | GTCAGAATTCTTATTACACCTTCCTCTTCTTCTTGGGCTGCAGCTTTTCGGCAGA |
| SUMO1 | TTCCACCTGAGCCGCCACTTCCACCTGAGCCGCC | AAGTGGCGGCTCAGGTGGAAGT |
| HMG 3-152 | ACGTGGTACCGCCATGGCAGGTGGTTCTGGTGGATCTCCTCTCCAAGAAAGTGAAG | GCGTTAGATATCCGCTCACTTGATCATCACCTT |
| HMG DBOX | GGGAGGAGGCTGCAGCAGCTCAACTAAGTCTTGAACAAG | AGCTGCTGCAGCCTCCTCCCTCTCTTCTTCTT |
| HMG HP1 | GGTGGAGGTGGATCTGGTGGAGGTGGATCTTCCACTGACGTTGATGAGCC | CCACCAGATCCACCTCCACCAATTTGGATTCCATGGTTGT |
| FRAG 3-152 | GACGAGCTGTACAAGGGTACCGGTGGTTCTGGTGGATCTGGTGGGGGCGGCGACCTG | AACCACCAGATCCACCAGAACCACCGAGAGGCAATTTGTCATGGAT |
| FRAG HP1 | TTCTGGTGGATCTGGTGGTTCTGGTGGATCTAATTCCTGTTCTGTGCGAC | TCCACCAGATCCACCGGTACCAGATCCACCAGAACCACCTCCTTCAGAGTCCTCACTG |
| SOD1 | CAAGGGTACCGGTGGTTCTGGTGGATCTATGGCGACGAAGGCCGTGTG | CACCGGTACCTTGGGCGATCCCAATTACAC |
| TRT | CAAGGGTACCGGTGGTTCTGGTGGATCTATGGCTTCTCATCGTCTGCT | CACCGGTACCTTCCTTGGGATTGGTGACGA |
| PML I | GGGAGACCCAAGCTGGCTAGCATGGTGAGCAAGGGCGAGG | TGCAGGCTCCATGGCGCTAGCAGAACCACCCTTGTACAGCTCGTCCATGCC |
| DAXX | GAGGCACGGTTGAAGCGTAA | CCATAGTCAGGGAAGGTATCAGG |
| ATRX | GGTCACTGCATGTAACAGCGT | GGGCACAATTAGTGCGGAATAA |

Table 3. siRNA information.

| Primer | species | Sequence |
|---|---|---|
| siSp100HMG-1 | homo sapiens | GGAATAACACCGCTGCAGCTGACAA |

| | | |
|---|---|---|
| si Sp100HMG-2 | homo sapiens | CCAGATGCTTCAGTCAAGTTCTCAG |
| siDAXX-1 | homo sapiens | GGAGUUGGAUCUCUCAGAA |
| siDAXX-2 | homo sapiens | GCCACACAATGCGATCCAGAA |
